# Multi-faceted roles of β-arrestins in G protein-coupled receptors endocytosis

**DOI:** 10.1101/2024.01.18.576020

**Authors:** Junke Liu, Li Xue, Magalie Ravier, Asuka Inoue, Julia Drube, Carsten Hoffmann, Eric Trinquet, Elodie Dupuis, Laurent Prézeau, Jean-Philippe Pin, Philippe Rondard

## Abstract

Internalization plays a crucial role in regulating the density of cell surface receptors and has been demonstrated to regulate intracellular signaling. Dysregulation of this process has been implicated in various diseases. The vast majority of GPCRs were considered to adopt one way for internalization. We challenged this conventional view by showing that multiple pathways converge to regulate the internalization of a specific receptor, based on an unparalleled characterization of 60 GPCR internalization profiles, both in the absence and presence of individual β-arrestins (βarrs). Furthermore, we revealed the internalization mechanism of the glucagon-like peptide-1 receptor (GLP-1R), a class B GPCR pivotal in promoting insulin secretion from pancreatic beta cells to maintain glucose homeostasis. GLP-1R can undergo agonist-induced internalization without βarrs, but can recruit and form stable complexes with βarrs. We found that GLP-1R recruits clathrin adaptor protein-2 for agonist-induced internalization in both βarr-dependent and -independent manners. These results provide a valuable resource for GPCR signaling and reveal the plasticity of different GPCRs to employ or not βarrs in the clathrin-mediated internalization.

## Introduction

Cell surface internalization of receptors and their trafficking into intracellular compartments are important for maintaining cell responses and homeostasis in a spatiotemporal manner^1–4^. Dysregulation of these processes has been implicated in various diseases^5^, such as obesity^6^, diabetes^7, 8^, and developmental and epileptic encephalopathy^9^.

G protein-coupled receptors (GPCRs) comprise the largest family receptors and regulate many kinds of intracellular signaling cascades in response to a diverse range of extracellular stimuli^10, 11^. To avoid overstimulation, these signals are terminated timely. Agonist stimulation leads to the phosphorylation of serine (Ser) and threonine (Thr) residues in the third intracellular loop (ICL3) and the carboxy-terminal domain (CTD) by GPCR kinases (GRKs) or the protein kinases ^12, 13^. The phosphorylated receptors recruit arrestins to trigger desensitization by blocking further G protein engagement, but also mediating the trafficking of activated receptors to clathrin-coated pits (CCPs) mainly through interacting with adaptor protein-2 (AP2) and clathrin^4, 14–16^. In addition to this arrestin-dependent way, some GPCRs are internalized through arrestin-independent pathways, such as fast endophilin-mediated pathway, and caveolae-dependent pathway^17^. Moreover, several GPCRs have been reported to be internalized constitutively (undergo internalization in the absence of agonist stimulation) in an arrestin-independent manner^18–20^. However, the internalization profiles of many GPCRs remain unclear and await exploration.

β-arrestin 1 (βarr1) and β-arrestin 2 (βarr2) are ubiquitously expressed arrestins and are believed to be major regulators for most G protein-coupled receptors (GPCRs) internalization. Although many GPCRs recruit βarrs, the GPCRs-βarrs interaction can differ between receptors, and various orientations of arrestins relative to the receptors were found by structure studies^21–25^. In addition, βarrs binds to GPCRs either through only the phosphorylated CTD (called tail-engaged) or through both the CTD and the transmembrane core of the GPCRs (called core-engaged)^26–28^. The core engagement is not necessary for receptor internalization^29^. Notably, among those GPCRs that recruit βarrs, someones need βarrs for internalization (eg. μ opioid receptor (MOR))^30^, but someones do not (eg. glucagon-like peptide-1 receptor (GLP-1R))^6, 8^. GLP-1R, a class B G protein–coupled receptor that promotes insulin secretion from pancreatic beta cells to regulate glucose homeostasis, is a key therapeutic target for type 2 diabetes (T2D) treatment^31^. Modulation of GLP-1R internalization improves agonist efficacy and achieves greater metabolic control in T2D with fewer side effects^8, 32^. However, the internalization mechanism of GLP-1R and whether βarrs play a role in this process remains elusive.

Based on different interaction modes of GPCRs and individual βarr, GPCRs historically have been grouped into two classes^16, 33, 34^. Type A receptors, which bind to βarr2 with higher affinity than βarr1, interact transiently with βarrs and rapidly recycle back from the endosomes to the plasma membrane. Type B receptors, which bind to βarr1 and βarr2 with equal affinity, form stable complexes with βarrs to stay longer time in endosomes, recycle back to the cell surface slowly, or be degraded by the lysosomes. Of note, distinct expression patterns of βarr1 and βarr2 across the human tissues were observed^35^, indicating the cellular background may also affect individual βarr‘s function. It was reported that the highly expressed βarr1 in MDA-MB231, facilitates cancer cell survival during hypoxia, but not βarr2^36^. Similarly, distinct contributions of individual βarr to various types of GPCR internalization were also found in HEK293 cells^37–41^. Hence, the individual βarr’s contribution to GPCR internalization remains debated and needs to be clarified systematically.

Here, by developing innovative time-resolved FRET assays and using βarrs knock-out HEK293 cells, we have analyzed the internalization profiles of 60 GPCRs. Most GPCRs exhibited βarr-independent constitutive internalization. Based on our agonist-induced internalization results, GPCRs can be classified into four groups: the receptors that cannot internalize, those that fully or partially require βarrs for internalization, and finally the ones that are βarrs independent for endocytosis. Furthermore, the GPCR CTD acts as a major molecular determinant for controlling distinct βarrs contribution to agonist-induced internalization. Finally, we have revealed several paths for GLP-1R agonist-induced internalization, in which the GRK-dependent clathrin-mediated path plays a major role. GLP-1R can recruit clathrin adaptor AP2 in βarrs-independent and -dependent manners by different regions at its CTD, and GRKs are required for these two processes. In addition, a GRK-independent path is also observed. These results suggest that the GPCR internalization process degenerates in biological systems, the ability of different elements to perform the same function^42^. Our studies provide unique data resources to dissect the contribution of individual βarr to GPCR internalization, opening an opportune pharmacological window for controlling GPCR signaling.

## Results

### Distinct expression patterns of βarrs in native tissues and cell lines

To investigate the contributions of βarrs to receptor internalization, we first need to quantify the level of βarr-1 and βarr-2 in the cells. Using the human proteome map database (www.Humanproteomemap.org)^35^, we show that the native expression level of βarr1 and βarr2 varies strongly across 30 different tissues and cell types **(Supplementary Figure 1)**. βarr1 is highly expressed in the retina and platelets, however, βarr2 is highly expressed in CD8 cells, NK cells, and monocytes, indicating that βarrs might have distinct functions in different types of cells. Next, we developed a time-resolved (TR) FRET-based assay to detect the endogenous expression of individual βarr in different cell lines. It is based on the binding of a pair of fluorescently labelled antibodies that recognize two non-overlapping epitopes at the cell surface of the native βarrs, and resulting in the measurement of TR-FRET signal **(Figure 1A, B)**. Interestingly, different expression patterns of βarrs in HEK293 cells and Hep G2 cells were observed. βarr1 is higher expressed in Hep G2 cells, whereas βarr2 is higher expressed in HEK293 cells and the background is measured with HEK-293 cell βarrs KO **(Figure 1A, B)**.

**Figure 1.**
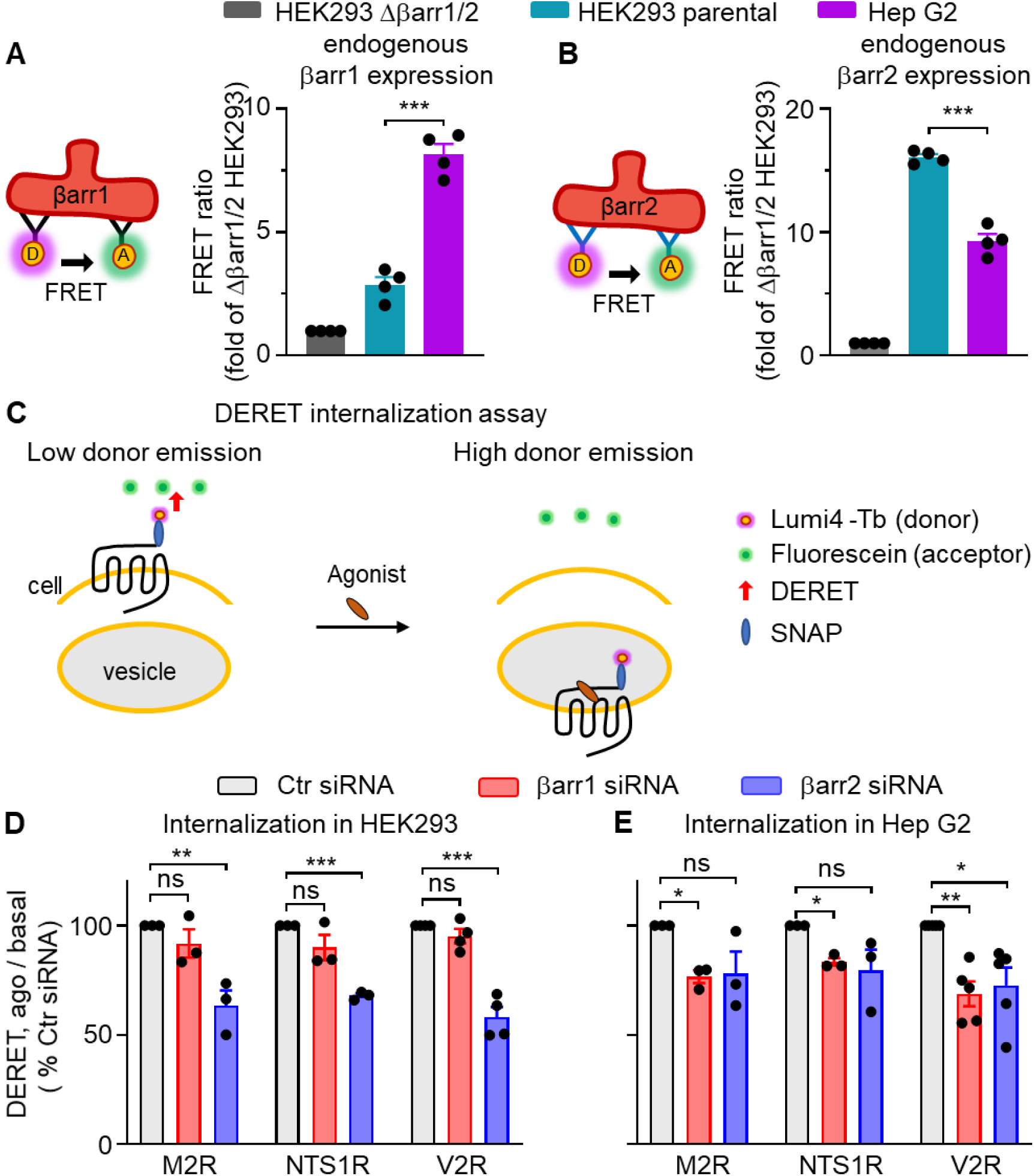
The contribution of the individual βarr to GPCR internalization depends on the cellular background. (A, B) Expression of endogenous βarr1 (A) or βarr2 (B) in parental HEK 293, Δβarr1/2 HEK 293 cells, and Hep G2 cells measured by a TR-FRET assay, where two antibodies highly specific for two distinct epitopes on βarr1 (A) or βarr2 (B), one labeled with a donor fluorophore and the other with an acceptor are used to generate a FRET signal. The total protein concentration is 1 mg/ml for each sample. Data are represented as mean ± SEM of four independent experiments performed in triplicates and normalized to Δβarr1/2 HEK 293 cells. Unpaired two-tailed t-test, ***p < 0.001. (C) Scheme illustrating the Diffusion-enhanced resonance energy transfer (DERET) internalization assay. SNAP-tagged receptors at the cell surface are labeled covalently with non-cell permeable SNAP-Lumi4®-Tb (energy donor, purple). When the receptor is at the cell surface, the addition of a free energy acceptor (fluorescein, green) to the cell medium leads to efficient energy transfer, quenching the donor signal and resulting in a low donor emission. Following agonist-induced internalization of the receptor, energy transfer to the acceptor is reduced, increasing the donor emission. (D, E) Internalization of M2R, NTS1R, and V2R with and without 60 min agonists stimulation (10 μM acetylcholine for M2R, 1 μM neurotensin for NTS1R, 1 μM vasopressin for V2R) in indicated cells transfected with control (Ctr) siRNA, βarr1 siRNA or βarr2 siRNA. Data are represented as mean ± SEM from 3-5 independent experiments performed in triplicates and normalized to Ctr siRNA. Unpaired two-tailed t-test, *p < 0.05. **p < 0.01, ***p < 0.001, ns, not significant.

### Contribution of the individual βarrs to GPCR internalization depends of cellular background

In these two cell lines, we show that the endogenous expression βarr1 or βarr2 affect differently the agonist-induced internalization of transfected GPCRs using βarr small interfering RNAs (siRNA). We investigate how amount of the endogenous βarrs in these two cell lines affect the agonist-induced internalization of transfected GPCRs by controlling the expression of βarr1 or βarr2 using small interfering RNAs (siRNA). The kinetics of internalization was monitored by the change of diffusion-enhanced resonance energy transfer (DERET) signal upon agonist stiumulation of the fluorescently labelled cell surface SNAP-tagged receptor (**Figure 1C**)^43^. Vasopressin V2 receptor (V2R), muscarinic acetylcholine receptor M2 (M2R) and neurotensin receptor type 1 (NTS1R), reported as βarr-dependent internalization^22, 34, 44, 45^, were used as models. In HEK293 cells, only the knockdown of βarr2 inhibited the agonist-induced internalization of these receptors (**Figure 1D**), showing that endogenous βarr2, but not βarr1, are involved in their internalization, and consistent with previous studies^37–41^. In contrast, in Hep G2 cells both βarr1 or βarr2 downregulation similarly inhibited the agonist-induced internalization of the receptors (**Figure 1D**), suggesting the involvement of both βarrs in these receptor internalization. As controls, a strong silencing of both βarrs expression (**Supplementary Figure 2**) and no or a slight effect on receptors cell surface expression (**Supplementary Figure 3**) were observed.

These data indicate that the expression level of the two βarrs is important for GPCR internalization. Indeed, the importance of βarr1 in GPCR internalization, especially when it is highly expressed, was undervalued in previous studies^37–41^.

### Importance of individual βarrs in GPCR internalization

To better determine the individual contribution of βarr1 and βarr2 to GPCR internalization in the absence of residual expression of βarrs in these knockdown approaches, we used the βarr1 and βarr2 double knockout (KO) HEK293 (Δβarr1/2) cells^38^. The effect of βarr1 and βarr2 in agonist-induced receptor internalization was then determined by transfecting these KO cells with optimized quantities of βarr1 or βarr2 cDNA, to obtain amounts of βarr1 or βarr2 much higher than their endogenous level measured in parental HEK-293 cells (**Supplementary Figure 4**).

The agonist-induced internalization of 60 unique transfected GPCRs transfected in Δβarr1/2 cells was then measured without and with co-expressed βarr1 or βarr2, using the DERET assay. The signal at 90 min after agonist stimulation, where the internalization was at its maximal for most GPCRs, was normalized to the basal signal of internalization (**Figure 2A-B**). The data revealed two main groups of GPCRs. First, 28 members did not exhibit agonist-induced internalization without βarrs, but the internalization of 19 of them was rescued by overexpression of either βarr1 (**Figure 2B**) or βarr2 (**Figure 2C**), showing that these receptors need βarrs for internalization. This group is called “totally βarr-dependent” for internalization (**Figure 2G**). The nine other receptors that did not show any evidence of internalization, even with βarr overexpression, are named “no internalization” (**Figure 2C-D, 2G**). For the second half of GPCRs (32 members) agonist-induced internalization in absence βarrs was detected (**Figure 2D**). For most of them, βarr1 or βarr2 overexpression increased internalization and they are named “partially βarr-dependent” for internalization (**Figure 2D, F, and G**). But interestingly, for some of them, the transfected βarr1 (12 receptors) or βarr2 (5) did not increase internalization, and they are named “βarr-independent” internalization (**Figure 2G**). Therefore, the 60 GPCRs are classified into four different groups (**Figure G**) with prototypical receptors and in good agreement with previous reports: (i) a minority of receptors are βarr-independent such PAR1^46, 47^ and vasoactive intestinal polypeptide type (VIP)-shared (VPAC1 and VPAC2) receptor^48, 49^ and more recently GLP-1R^6, 8^; (ii) most of them are partially βarr-dependent such as the vasopressin V1a (V1AR) and V2 (V2R) receptors, and β_2_ adrenergic receptor (β_2_AR); and (iii) totally βarr-dependent such as the μ-opioid (MOR) and M2 receptors; and (iv) finally, 15% of the receptors analyzed (among these half are from class C) did not show any evidence of internalization in Δβarr1/2 KO cells, consistent previous data showing that they not recruit βarr upon agonist stimulation, such as AT2^50^, mGluRs^51, 52^ and GABA_B_ receptor^52^. Similar results were obtained for both βarrs, consistent with βarr1 or βarr2 which can play a similar role in internalization, and consistent with our data of βarrs knockdown in Hep G2 cells (**Figure 1D**). However, for some receptors, βarr2 has a stronger effect than βarr1 in promoting receptor internalization (**Supplementary Figure 5**). It could explain the lower proportion of “βarr independent” receptors with βarr2 compared to βarr1 conditions (**Figure 2G**). Altogether, these findings show that the βarrs have a distinct role in the agonist-induced internalization of the different receptors.

**Figure 2.**
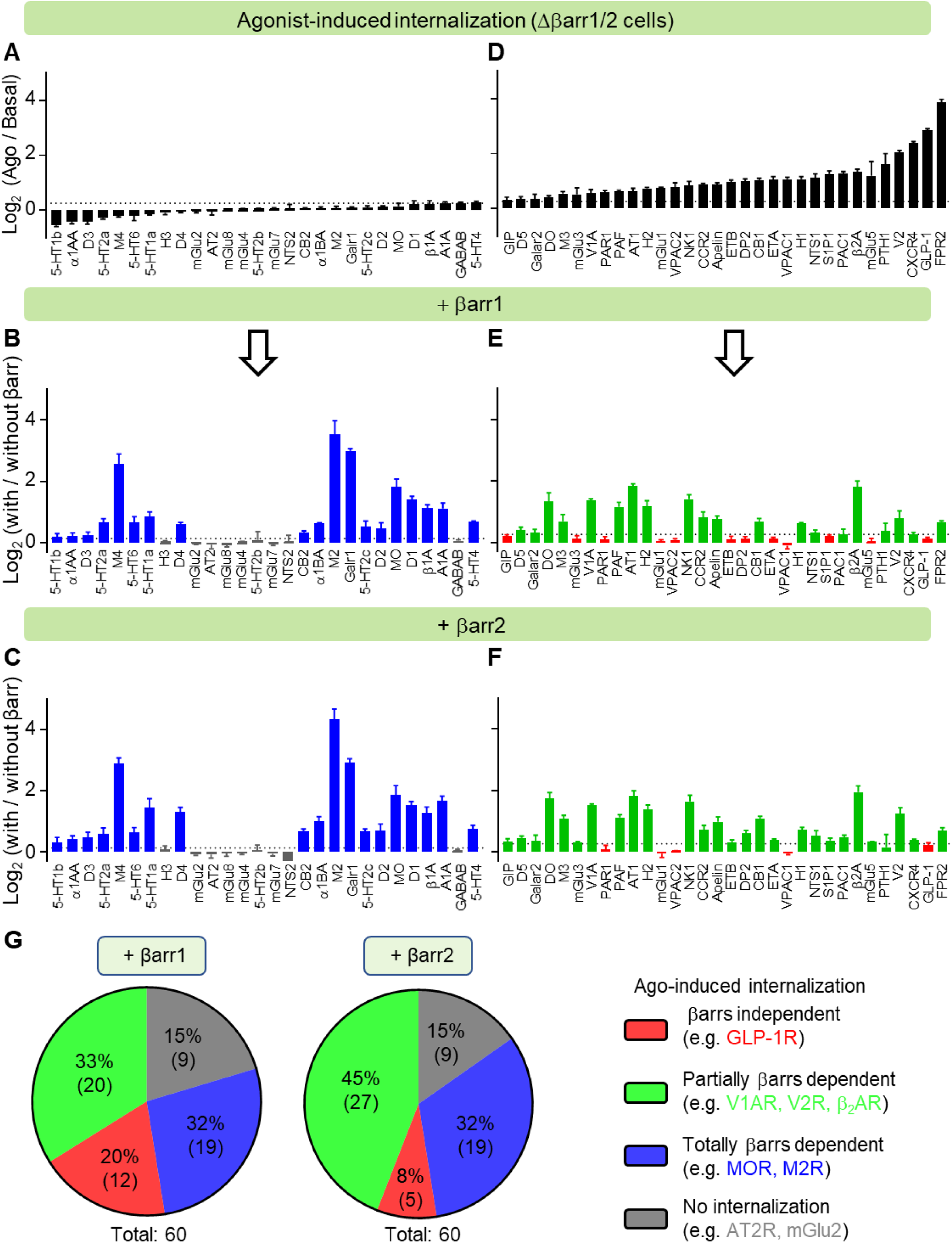
Distinct effects of βarrs on agonist-induced internalization for 60 GPCRs. (A, D) Internalization of 60 GPCRs with and without 90 min agonist stimulation in Δβarr1/2 HEK 293 cells. (B, E) Effect of transfected βarr1 on the agonist-induced internalization of 60 GPCRs in Δβarr1/2 HEK 293 cells. (C, F) Effect of transfected βarr2 on the agonist-induced internalization of 60 GPCRs in Δβarr1/2 HEK 293 cells. (G) Classification of 60 GPCRs depends on the effect of either βarr1 or βarr2 on the agonist-induced internalization. (A, B, C, D, E, F) Data are represented as mean ± SEM from 3-4 independent experiments performed in triplicates.

We also measured the constitutive internalization of these 60 GPCRs in Δβarr1/2 cells. All of these GPCRs exhibited constitutive internalization (**Supplementary Figure 6**), except the serotonin 5-HT2c and dopamine D5 receptors. And βarr1 or βarr2 overexpression had no or only a slight effect the constitutive internalization. These data indicate that constitutive internalization of the different GPCRs maybe share a general mechanism which is independent of βarrs.

### βarrs are not required for GLP-1R internalization

We further analyzed the internalization of GLP-1R that was βarr-independent^6^ and we used as control the MOR that has a totally dependent internalization^30^. The kinetics of GLP-1R internalization was not changed by re-introducing βarr1 or βarr2 in Δβarr1/2 cells (**Figure 3A**), and internalization was independent of the agonist concentration. In contrast, agonist-induced internalization of MOR was rescued by transfected βarrs (**Figure 3B**). We verified that the experiments were performed at a similar cell surface expression level of two receptors (**Supplementary Figure 7**). Consistently, the agonist-induced internalization of transfected GLP-1R in parental HEK293 cells was not affected by the cotransfected of either βarr1 or βarr2 (**Figure 3B and Supplementary Figure 8**). This is in contrast to the MOR where the reduced expression lower expression level of βarr2 impaired agonist-induced internalization (**Figure 3D and Supplementary Figure 8A**). As a control, we verified that either βarr1 or βarr2 or both (**Supplementary Figure 8B-C**) was strongly downregulated and the experiments were performed under similar amounts of cell surface GLP-1R and MOR (**Supplementary Figure 8D**). These data further demonstrate that βarrs are not required for GLP-1R internalization, but they are necessary for MOR.

**Figure 3.**
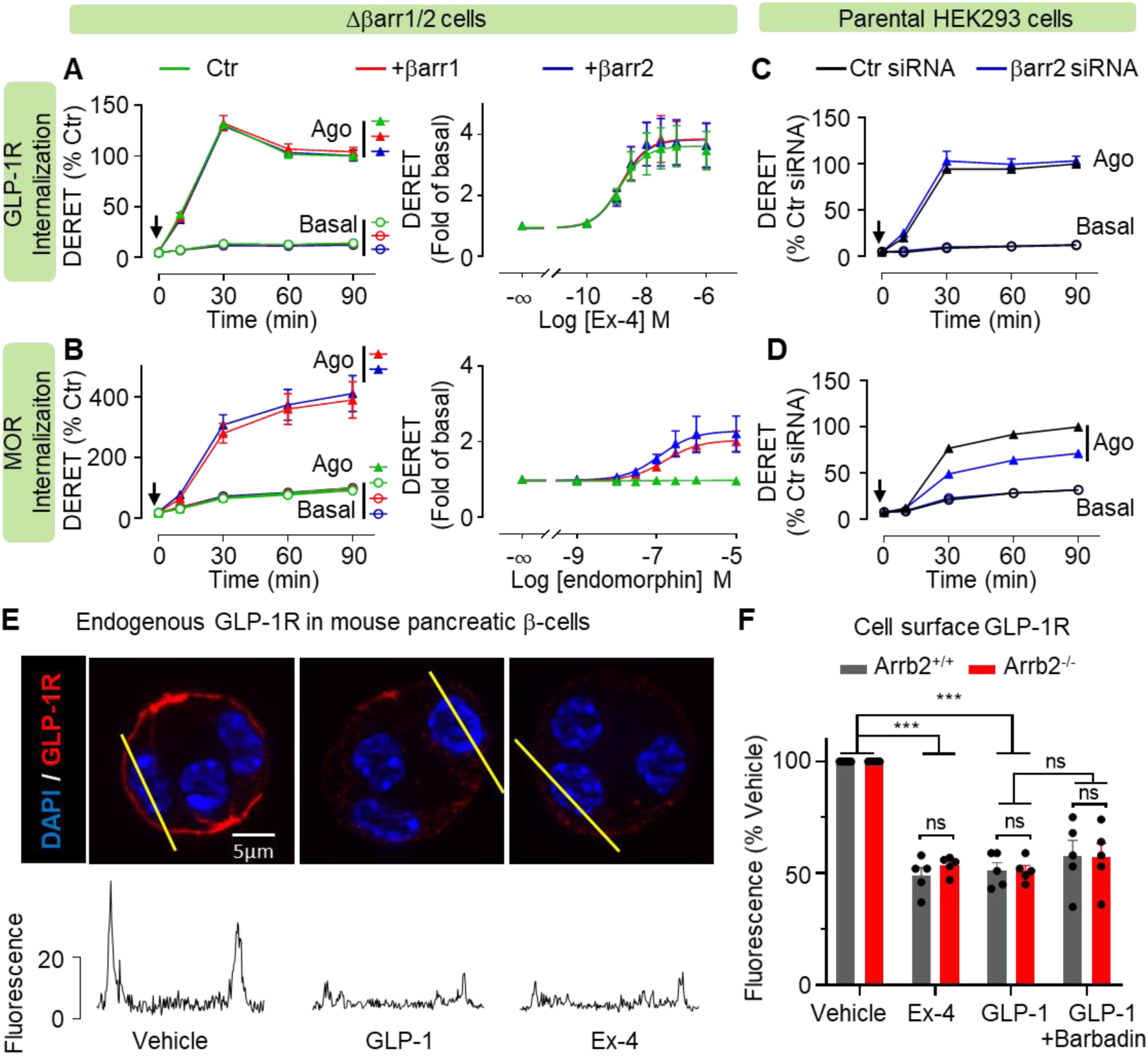
βarrs are not required for GLP-1R internalization. (A, B) Internalization of GLP-1R and MOR with and without agonist stimulation in Δβarr1/2 HEK 293 cells. Left, the kinetic curves (0.1 μM exendin-4 for GLP-1R, 2 μM endomorphin-1 for MOR). Right, the dose-response curve following 60 min agonists stimulation. (C, D) Internalization of GLP-1R and MOR with and without agonist (0.1 μM exendin-4 for GLP-1R, 2 μM endomorphin-1 for MOR) stimulation in parental HEK 293 cells transfected with indicated siRNA. (E) Images of endogenous cell surface expressing GLP-1R (Red) with and without 25 min agonists stimulation (10 nM GLP-1, 100 nM exendin-4) in mouse pancreatic β-cells. Nuclei are stained by DAPI (Blue). Images are from one representative experiment of 5 independent experiments. Scale bar 5 μm. (F) Quantification of endogenous cell surface GLP-1R reduction with and without 25 min exendin-4 (100 nM), GLP-1 (10 nM), and GLP-1 (10 nM) with barbadin (30 μM) stimulation in mouse pancreatic β-cells. Data are represented as mean ± SEM from 5 independent experiments. Two-way ANOVA test, ***p < 0.001, ns, not significant.

We then analyzed the internalization of endogenous GLP-1R in mouse pancreatic β-cells. Agonists GLP-1 and Exendin-4 decreased the amount of cell surface receptors detected with a fluorescent anti-GLP-1R antibody in pancreatic β-cells derived from WT mice (**Figure 3E**). Interestingly, similar results were obtained with pancreatic β-cells from βarr2 KO mice (**Figure 3F**). To rule the possibility that βarr1 favors GLP-1R internalization in these cells, we show that barbadin, a selective blocker of AP2-βarr interaction that inhibit the internalization of GPCRs^44^, has no effect (**Figure 3F**). Altogether, our data show that βarrs are not required for endogenous GLP-1R internalization in pancreatic β-cells.

We verified that the βarr-independent internalization of GLP-1R was not due to its lack of interaction with the βarrs. First, we have developed a TR-FRET assay to monitor the recruitment of the endogenous βarr2 to the fluorescent GLP-1R tagged at the C-terminal end with a GFP (**Figure 4A**). After agonist stimulation, a FRET signal is measured between an antibody linked to a fluorophore donor that recognizes the endogenous βarr2 and the GFP used as an acceptor. In these experiments, both GLP-1R and MOR produced a strong increase of FRET signal upon agonist stimulation (**Figure 4A**) and this signal was further increased when βarr2 was transfected. Second, we also analyzed the ability of GLP-1R to induce the formation of complexes between βarrs and AP2 which is an important initial step to stabilize GPCRs into CCPs for internalization^16^. We then developed a TR-FRET assay to detect the proximity between the endogenous βarr2 and AP2 upon the agonist-induced GLP-1R and MOR activation (**Figure 4B**). This assay is based on the use of two antibodies specific to βarr2 and AP2 and labeled with a donor and an acceptor fluorophore, respectively. Both GLP-1R and MOR were able to induce the formation of the endogenous complex between βarr2 and AP2 (**Figure 4B**).

**Figure 4.**
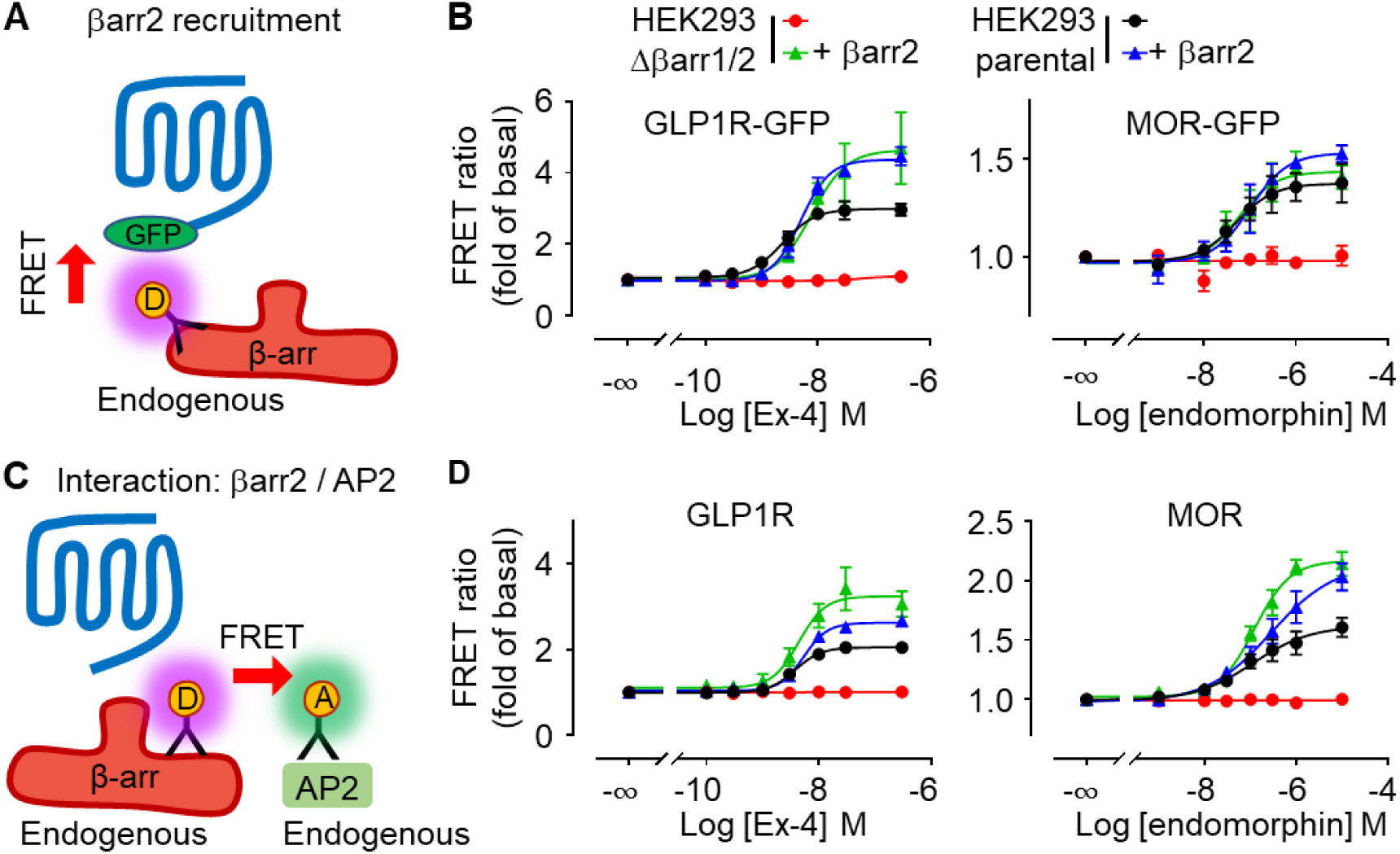
GLP-1R recruits βarrs and induces βarrs/AP2 complex formation. (A) Scheme illustrating measuring βarr2 recruitment by a TR-FRET assay, where one antibody highly specific for βarr2 labeled with a donor fluorophore and the receptors fused with an EGFP as an acceptor are used to generate a FRET signal. (B) Dose-response curves of βarr2 recruitment by GLP-1R (left) and MOR (right) in indicated cells with and without overexpression βarr2. (C) Scheme illustrating measuring βarr2 and AP2 interaction by a TR-FRET assay, where one antibody highly specific for βarr2 labeled with a donor fluorophore and the other highly specific for AP2 labeled with an acceptor fluorophore are used to generate a FRET signal. (B) Dose-response curves of βarr2 recruitment by GLP-1R (left) and MOR (right) in indicated cells with and without overexpression βarr2. (D) Dose-response curves of βarr2 and AP2 interaction induced by GLP-1R (left) and MOR (right) in indicated cells with and without overexpression βarr2. Data are represented as mean ± SEM from 3-4 independent experiments.

Altogether, these results show that GLP-1R can induce the formation of βarrs-AP2 indicating that βarrs could be involved in GLP-1R internalization, even though they are not required for its internalization.

### The C-terminal region of GLP-1R is responsible for βarr-independent internalization

We then aimed to identify the molecular determinants in GLP-1R that are responsible for internalization in the absence of βarrs. When the C-terminal region (CTD) of GLP-1R is replaced by that of MOR (**Figure 5A**), in the chimera GLP-1R_MOR_, the receptor becomes βarr partially dependent for internalization in Δβarr1/2 cells (**Figure 5B**), similarly to the B2AR, V1A and V2 receptors (**Supplementary Figure 5**). Indeed, overexpression of βarr-1 or -2 improved strongly internalization (**Figure 5B**, **Supplementary Figure 10A and 11**).

**Figure 5.**
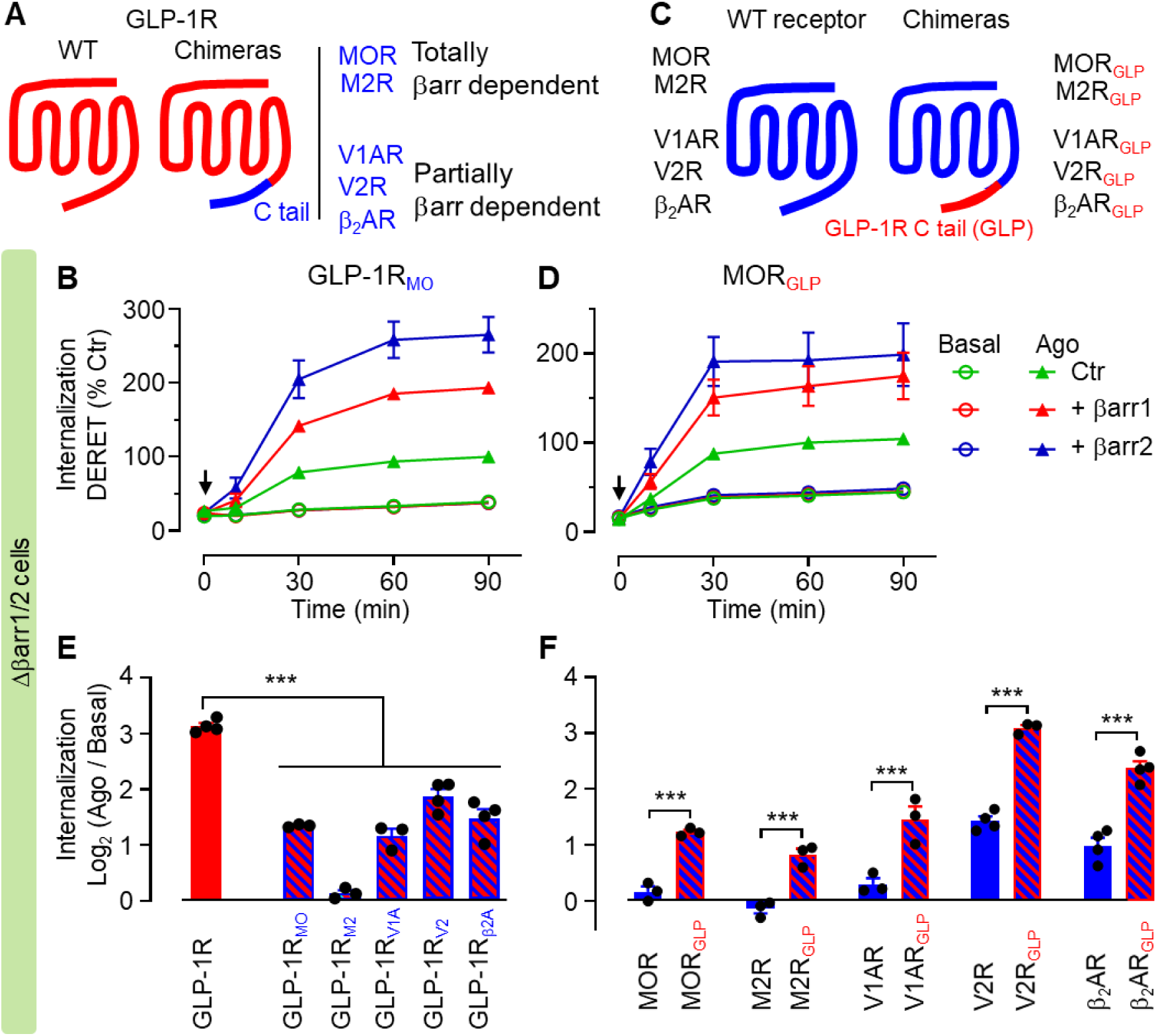
The C-terminal region of GLP-1R is responsible for βarr-independent internalization. (A, C) Scheme illustrating the CTD exchanged chimeras for βarr independent receptor (GLP-1R) and totally βarr dependent receptors (MOR, M2R), partially βarr dependent receptors (V1AR, V2R, β_2_AR). (B) Internalization of chimeric receptors (GLP-1R_MO_, GLP-1R with MOR CTD; MOR_GLP_, MOR with GLP-1R CTD) with and without agonists stimulation in Δβarr1/2 HEK 293 cells. (E, F) Internalization of chimeric receptors with and without 60 min agonists stimulation in Δβarr1/2 HEK 293 cells. 0.1 μM exendin-4 is used for WT GLP-1R and the chimeric GLP-1R with the CTDs from the other receptors. 2 μM endomorphin-1 is used for MOR and MOR_GLP_. 10 μM acetylcholine is used for M2R and M2R_GLP_. 1 μM vasopressin is used for V1AR, V2R, V1AR_GLP_, and V2R_GLP_. 10 μM isoproterenol is used for β_2_AR and β_2_AR_GLP_. Data are represented as mean ± SEM from 3-4 independent experiments.

Conversely, the reverse chimera made of MOR with the CTD of GLP-1R (**Figure 5C**), MOR_GLP_, becomes partially βarr-independent for internalization (**Figure 5D, Supplementary Figure 10, 11**), even though overexpression of βarr-1 or -2 improved internalization in Δβarr1/2 cells (**Supplementary Figure 10C**). Similar results were obtained with GLP-1R chimera made of the CTD of other receptors “totally βarr-dependent” (M2) and “partially βarr-dependent” (V1AR, V2R, and β_2_AR) for internalization (**Figure 5E**). In addition, similarly to the chimera MOR_GLP_, the CTD of GLP-1R changed the behavior of the receptors above that are βarr-dependent for internalization, since their chimera became βarr-independent for internalization (**Figure 5F)**. We have shown that overexpression βarr-1 or -2 improved the agonist-induced internalization of all the chimera except the ones made between the “partially βarr-dependent”, V1A_GLP_, V2R_GLP_, and β2AR_GLP_ internalization, the CTD of GLP-1R being sufficient for an optimal agonist-induced internalization even in the absence of βarr (**Supplementary Figure 10A-C**). We have controlled that the cell surface expression of the SNAP chimera was not affected by the co-expression of the βarrs (**Supplementary Figure 11**). Altogether, our results show that the CTDs of GPCRs are the major determinant for controlling the contribution of βarr to internalization.

### Key determinants in the proximal CTD of GLP-1R for internalization

Among the receptors tested above, M2R has the shortest CTD (**Supplementary Figure 13**) and GLP-1R_M2_ showed no agonist-induced internalization without βarrs (**Figure 5E**), indicating that the CTD is important for GLP-1R internalization^53^. To identify the key determinants in GLP-1R CTD, we analyzed deletion mutants (Δ431, Δ440, Δ446, and Δ451) (**Figure 6A**) that were all correctly expressed at the cell surface (**Figure 6B**). The shortest CTD constructs (Δ431 and Δ440) strongly reduced agonist-induced internalization, whereas no effects were measured for the longest CTD (Δ446 and Δ451) (**Figure 6C**). This demonstrated that residues 440-445 are most likely to be involved in GLP-1R internalization. Consistent with this, we observed substantially reduced agonist-induced internalization by mutating residues 440-445 (*6A*) and 438-445 (*8A*) to Ala, respectively (**Figure 6D**). Of note, the mutant C438A, which is no longer palmitoylated following exendin-4 stimulation (REF), internalizes similarly to the wild-type GLP-1R (**Figure 6D**), and is not in agreement with previous report^6^. As a control, we have shown that the mutations of the four phosphorylable residues in the distal part of the CTD, Ser451, Ser452, Thr455 and Thr457^21^ in the mutant *2A* (S451A, S452A) and *2A* (S451A, S452A, T455A, and T457A) did not affect internalization (**Figure 6D**). We also verified that all mutants are correctly expressed at the cell surface (**Figure 6B**). Overall, these data demonstrate that residues 440-445 are important for GLP-1R internalization.

**Figure 6.**
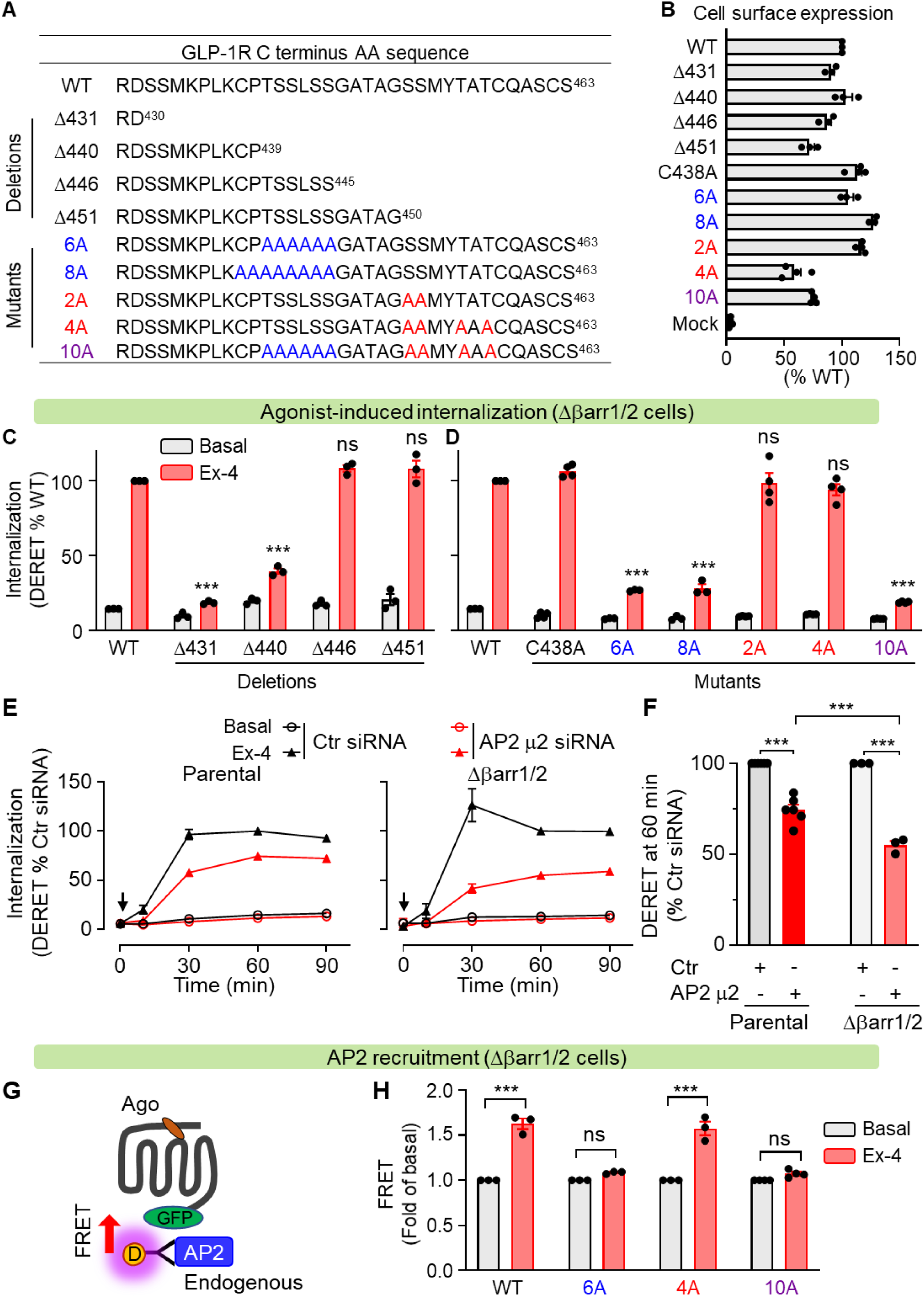
AP2-mediated the agonist-induced internalization of GLP-1R. (A) Scheme illustrating the CTD deletions and mutants for GLP-1R. (B) Cell surface expression by SNAP-tag labeling assay of the GLP-1R deletions and mutants. (C, D) Internalization of GLP-1R WT and indicated mutants with and without 60 min agonist (0.1 μM exendin-4) stimulation in Δβarr1/2 HEK 293 cells. (E, F) Internalization of GLP-1R with and without 60 min agonist (0.1 μM exendin-4) stimulation in indicated cells transfected with control (Ctr) siRNA or AP2 μ2 siRNA. (G) Scheme illustrating measuring AP2 recruitment by a TR-FRET assay, where one antibody highly specific for AP2 labeled with a donor fluorophore and the receptors fused with an EGFP as an acceptor are used to generate a FRET signal. (H) AP2 recruitment by WT GLP-1R and indicated mutants in Δβarr1/2 HEK 293 cells. Data are represented as mean ± SEM from 3-6 independent experiments. Unpaired two-tailed t-test, ***p < 0.001, ns, not significant.

### AP2 mediates the agonist-induced internalization of GLP-1R

Next, we aimed to delineate the mechanism responsible for GLP-1R internalization in Δβarr1/2 cells, by using RNAi screening. It was reported that caveolin-1 regulates the trafficking of GLP-1R^54^. However, we did not observe any effect of either caveolin-1, caveolin-2, or caveolin-3 siRNA on agonist-induced GLP-1R internalization both in HEK293 parental cells and Δβarr1/2 cells (**Supplementary Figure 13**). In addition, we did not observe any effect of small GTPase ARF6 siRNA (**Supplementary Figure 13**), which assists the recruitment of AP2 and clathrin to βarrs for regulating receptor internalization^55^.

Interestingly, we have shown that AP2 is necessary for agonist-induced GLP-1R internalization. Indeed, it was strongly inhibited in AP2 μ2 siRNA-transfected HEK293 parental and Δβarr1/2 cells (**Figure 6E-F**) for a similar downregulation of the expression of the AP2 μ2 subunit (**Supplementary Figure 14A**). But this inhibitory effect is stronger in Δβarr1/2 cells compared to that in parental cells which suggests that βarrs might be involved in GLP-1R internalization in parental cells. Consistent with this, in these cells GLP-1R agonist-induced internalization was inhibited stronger by transfecting a combination of AP2 μ2 and βarr2 siRNA than with individual transfection (**Supplementary Figure 14B-C**).

### GLP-1R recruits AP2 by its proximal CTD and independently of βarrs

To examine how GLP-1R recruits and interacts with AP2, we developed a TR-FRET assay that measures the proximity between one antibody, labeled with a donor fluorophore, that binds the endogenous AP2, and the C-terminal tagged GFP GLP-1R where the GFP is used as an acceptor fluorophore (**Figure 6G**). The agonist induced a strong increase of FRET signal in Δβarr1/2 (**Figure 6H**), which is not measured for the GLP-1R *6A* mutant in the proximal part of the CTD while, but observed for the *4A* mutant in the distal part of the CTD (**Figure 6H**). These data show that GLP-1R recruits AP2 during agonist-induced internalization, independently of βarrs, and that the residues 440-445 in the CTD domain play an important role in this process.

### βarrs favor AP2 recruitment to GLP-1R

Next, we addressed how βarrs are involved in the AP2 recruitment of GLP-1R since GLP-1R recruits βarrs **(Figure 4A)** and can induce the formation of a direct interaction between βarrs and AP2 **(Figure 4B)**. As observed in Figure 6H, GLP-1R recruited strongly AP2 in the Δβarr1/2 cells in the absence of βarrs, even though overexpressed βarr2 largely improved this recruitment (**Figure 7A**). The importance of βarrs in facilitating AP2-mediated internalization is illustrated by the mutant *6A* strongly affecting receptor internalization in the Δβarr1/2 cells due to mutations in the AP2 binding site (**Figure 7A, 7B**), but partially restored upon βarr-2 overexpression. As a control, we show that mutant *6A* can induce the formation of a complex between βarr and AP2 (**Figure 7E**). The importance of βarrs is also shown by the mutant *4A* (**Figure 7A, 7B**) which is no longer sensitive to this overexpressed βarr2 since it is not able to recruit βarr2 (**Figure 7D**). As a control, mutation of both AP2 and βarr binding sites in the mutant *10A* abolished AP2 recruitment (**Figure 7A**) and βarr (**Figure 7D**) and no evidence of βarr2-AP2 complex was observed (**Figure 7E**). For example, the MOR-GFP construct required the presence of βarr to measure AP2 recruitment in the Δβarr1/2 cells (**Supplementary Figure 15A**). It is consistent with the GLP-1R

**Figure 7.**
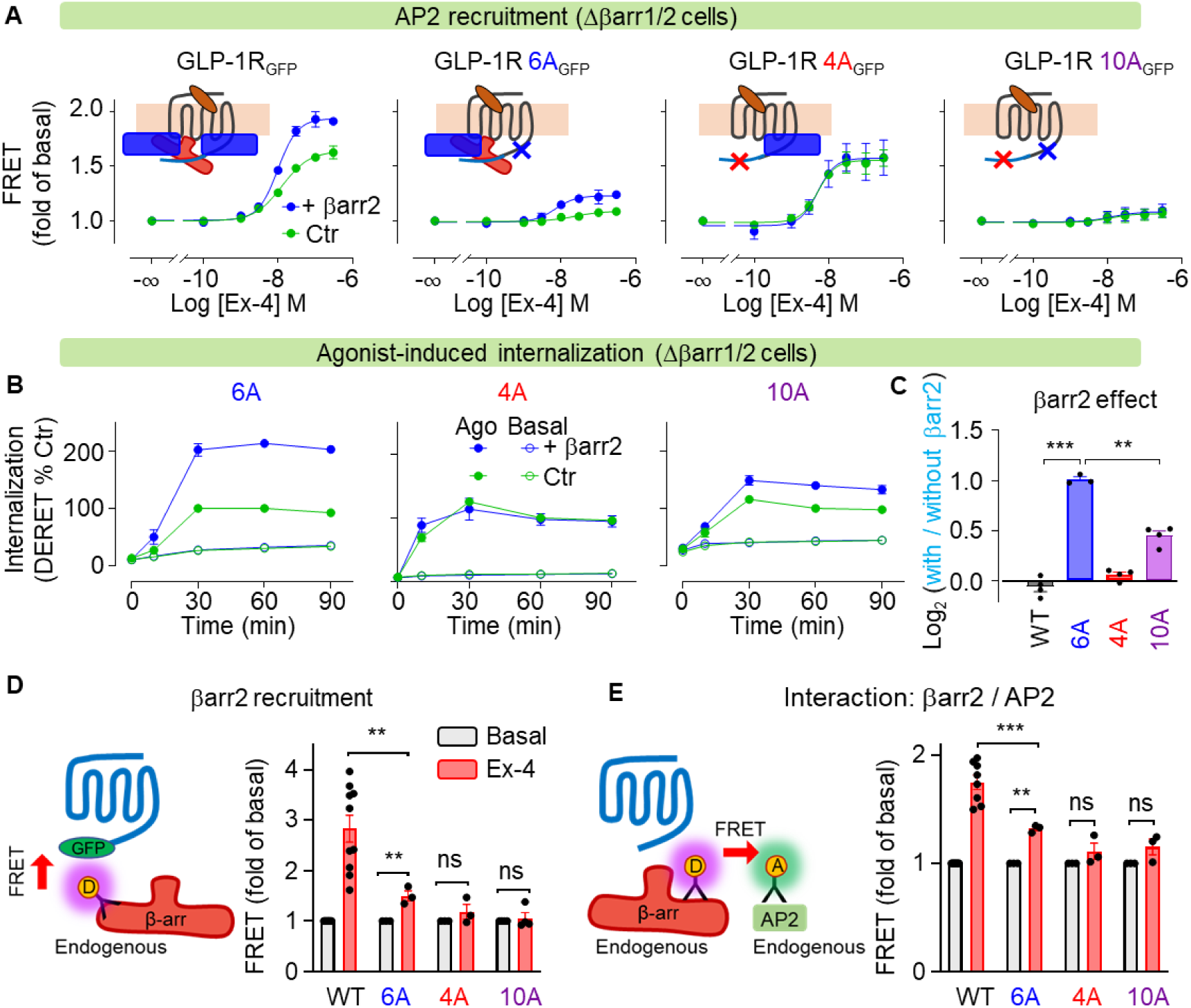
βarrs are involved in the AP2 recruitment and agonist-induced internalization of GLP-1R. (A) Dose-response curves of AP2 recruitment by GLP-1R WT and indicated mutants in Δβarr1/2 HEK 293 cells transfected with and without βarr2. (B) Internalization of GLP-1R and indicated mutants with and without agonist (0.1 μM exendin-4) stimulation in Δβarr1/2 HEK 293 cells transfected with and without βarr2. (C) Effect of transfected βarr2 on the agonist-induced internalization of GLP-1R and indicated mutants in Δβarr1/2 HEK 293 cells. (D) βarr2 recruitment by GLP-1R and indicated mutants with and without agonist (0.1 μM exendin-4) stimulation in Δβarr1/2 HEK 293 cells transfected with βarr2. (E) βarr2 and AP2 interaction induced by GLP-1R and indicated mutants with and without agonist (0.1 μM exendin-4) stimulation in Δβarr1/2 HEK 293 cells transfected with βarr2. Data are represented as mean ± SEM from 3-4 independent experiments. Unpaired two-tailed t-test, **p < 0.01, ***p < 0.001, ns, not significant.

Altogether, our results are consistent with a model where GLP-1R can recruit AP2 in both βarr-independent and dependent manners (**Figure 7A**), with most probably a high dominance of the βarr-independent mechanism since AP2 seems sufficient to induce GLP-1R internalization. The phosphorylable sequence 440-TSSLSS-445 is involved in the AP2 recruitment, while the motif 451-SSMYTAT-457 is responsible for the βarr recruitment.

Overall, these data demonstrate that GLP-1R was internalized via βarr-independent and dependent ways. The independent way was the main process, whereas the dependent way may act as a backup role, which can be observed when the main process is blocked (**Figure 7B**). Moreover, these data demonstrate that GLP-1R recruits AP-2 in both βarrs-dependent and independent manners by distinct regions at the CTD, indicating a tripartite interaction exists between GLP-1R, βarrs, and AP2 (each protein involved in a direct interaction with the other two proteins). Interestingly, each interaction is independent of the others but cooperates. βarrs promote GLP-1R to recruit AP2 (**Figure 7A**), on the other way AP2 also facilitates βarrs recruitment. As βarrs recruitment was impaired by the 6A mutant (**Figure 7D**).

### GRK-mediated AP2 and βarrs recruitment during GLP-1R internalization

We then asked whether GRKs are needed for βarrs and AP2 recruitment, and the internalization of GLP-1R by using HEK293 cells lacking GRK2/3/5/6 (ΔGRK2/3/5/6 cells)^15, 45^. As observed in Figure 8A-B, AP2 and βarr2 recruitment were completely abolished in ΔGRK2/3/5/6 cells, and were rescued by re-introducing each individual GRK. Similarly, the agonist-induced internalization was substantially reduced without GRK2/3/5/6 (**Figure 8C, Supplementary Figure 17A, B**). These results suggest that GRKs involved in AP2 and βarrs recruitment during GLP-1R internalization. Notably, we still observed agonist-induced GLP-1R internalization in the absence of GRK2/3/5/6 (**Figure 8C, Supplementary 18B**), indicating a GRK-independent way for GLP-1R internalization. In contrast, we only observed GRK-dependent internalization for MOR^45^ (**Supplementary Figure 17C**).

**Figure 8.**
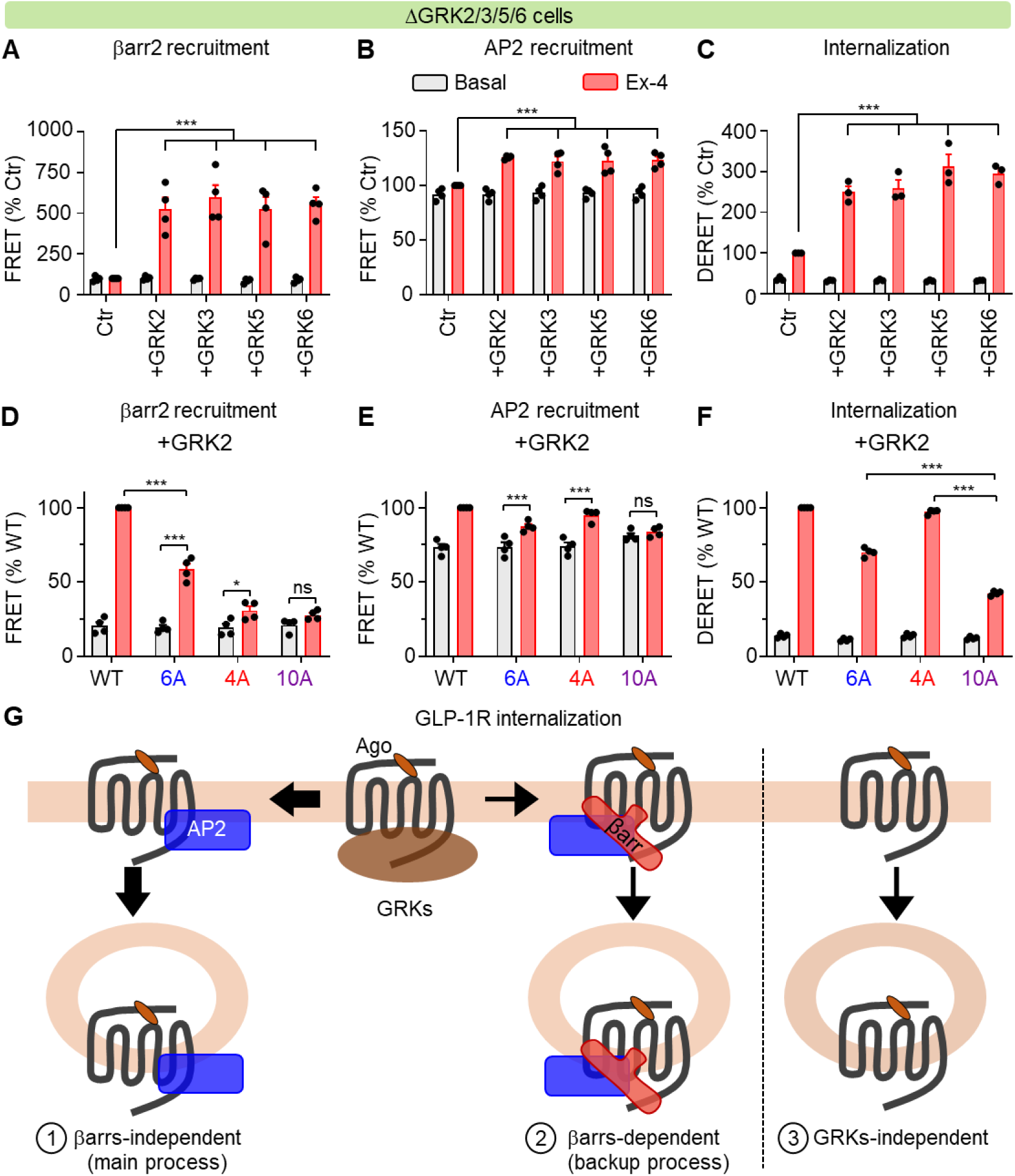
GRKs-mediated βarrs recruitment, AP2 recruitment, and the agonist-induced internalization of GLP-1R. (A, D) βarr2 recruitment by GLP-1R WT and indicated mutants with and without agonist (0.1 μM exendin-4) stimulation in ΔGRK2/3/5/6 HEK 293 cells transfected with and without indicated individual GRKs. (B, E) AP2 recruitment by GLP-1R WT and indicated mutants with and without agonist (0.1 μM exendin-4) stimulation in ΔGRK2/3/5/6 HEK 293 cells transfected with and without indicated individual GRKs. (C, F) Internalization of GLP-1R WT and indicated mutants with and without 60 min agonist (0.1 μM exendin-4) stimulation in ΔGRK2/3/5/6 HEK 293 cells transfected with and without indicated individual GRKs. (A, B, C, D, E, F) Data are represented as mean ± SEM from 3-4 independent experiments. Unpaired two-tailed t-test, *p < 0.05. **p < 0.01, ***p < 0.001, ns, not significant. (G) Schematic representation of the mechanism governing GLP-1R internalization. Multiple pathways are implicated in the regulation of GLP-1R internalization. GLP-1R engages AP2 in both barr-independent (Main process) and barr-dependent (backup process) manners for agonist-induced internalization, with GRKs playing a pivotal role in both processes. Furthermore, an alternative GRK- independent pathway has been identified.

To determine the roles of GRKs in the recruitment of AP2 and agonist-induced internalization of GLP-1R in βarrs-dependent and -independent manners, the 4A, 6A, and 10A mutants were examined. The results showed that the 10A mutant completely abolished the effect of GRK2/3/5/6 on AP2 recruitment (**Figure 8E, Supplementary Figure 18**), leading to lower agonist-induced internalization compared to the 4A and 6A mutants (**Figure 8F, Supplementary Figure 18**). These indicate that GRKs are required for both βarrs-dependent and -independent AP2 recruitment and internalization of GLP-1R. However, the 10A mutant did not completely block agonist-induced internalization (**Figure 8F, Supplementary Figure 18**), suggesting the existence of a GRK-independent pathway for GLP-1R internalization.

Overall, these data demonstrate that various internalization pathways have evolved to regulate GLP-1R internalization (**Figure 8G**). GLP-1R recruits AP-2 in both βarr-dependent and independent manners for agonist-induced internalization, and GRKs are essential for these two processes. In addition, a GRK-independent way is also observed (**Figure 8G**).

### Internalization of GLP-1R genetic variants in AP2 binding motif is favored by βarrs

Natural genetic variation are highly relevant for diseases^51, 56^. We show that two of the three natural genetic GLP-1R variants^56^, T440A, L443P, and S445T, located in the AP2 binding motif **(Figure 9A)**, affect agonist-induced internalization. Indeed, two mutants where phosphorylable residues are changed, T440A and S445T, have a significantly reduced agonist-induced internalization in Δβarr1/2 cells **(Figure 9B)**, an effect that is rescued by overexpression of βarr1 **(Supp Figure 20A)** or βarr2 (**Figure 9B**). As controls, the mutant L443P did not show a significant effect **(Figure 9C)**, and a similar expression surface of the ^ST^GLP-1R constructs was measured **(Supp Figure 20B)**. Consistently, we show that the AP-2 recruitment is significantly impaired for these two mutants T440A and S445T with an impaired internalization **(Figure 9E)**, but no effect is observed for the other one. In addition, none of them impaired significantly βarr recruitment **(Figure 9F)**, consistent with the fact that the βarr binding motif is not impaired.

**Figure 9.**
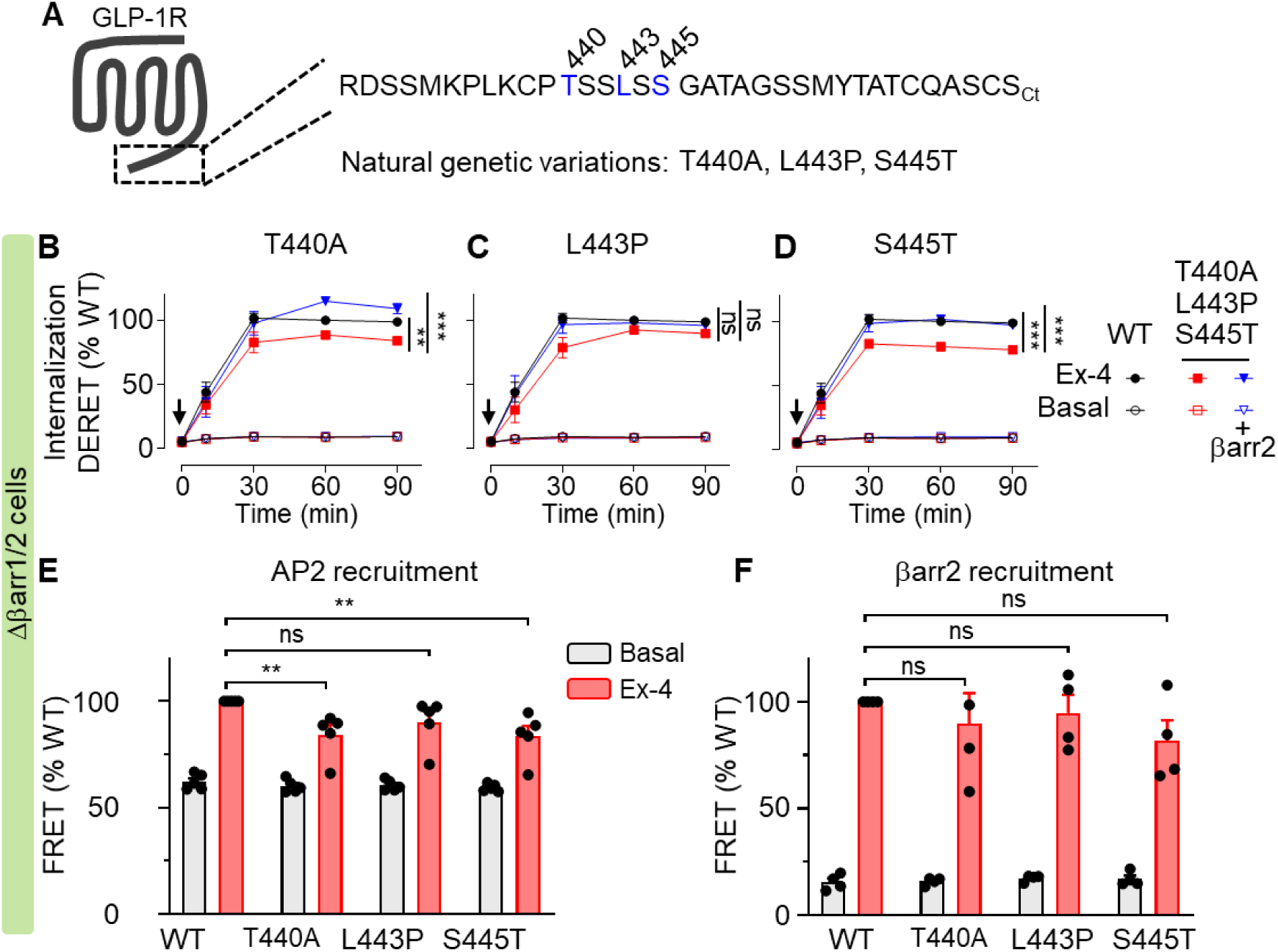
Natural genetic variants of GLP-1R partially require βarrs for agonist-induced internalization. (A) Scheme illustrating GLP-1R genetic variants in AP2 binding motif. (B, C, D) Internalization of GLP-1R and indicated variants with and without agonist (0.1 μM exendin-4) stimulation in Δβarr1/2 HEK 293 cells transfected with and without βarr2. (E) AP2 recruitment by WT GLP-1R and indicated variants with and without agonist (0.1 μM exendin-4) stimulation in Δβarr1/2 HEK 293 cells. (F) βarr2 recruitment by GLP-1R and indicated variants with and without agonist (0.1 μM exendin-4) stimulation in Δβarr1/2 HEK 293 cells transfected with βarr2. (B, C, D, E, F) Data are represented as mean ± SEM from 3-5 independent experiments. Unpaired two-tailed t-test, *p < 0.05. **p < 0.01, ***p < 0.001, ns, not significant.

## Discussion

GPCRs are targeted by more than 30% of approved drugs, modulating their internalization processes may maximize therapeutic efficacy and minimize adverse envents^8, 57^. Here, we provide an unprecedented description of 60 GPCR internalization profiles, including both constitutive and agonist-induced internalization in the absence and presence of βarrs. Most GPCRs may share a general mechanism for constitutive internalization, which is independent of βarrs (Supplementary Figure), or clathrin^18^. In contrast, the agonist-induced process is much more diverse. By dissecting βarrs contribution to the agonist-induced internalization, we classified these 60 GPCRs into four groups: (1) the receptors that cannot internalize, those (2) fully or (3) partially require βarrs for internalization, and finally (4) the ones that are βarrs independent for endocytosis (**Figure 2**). This dataset of internalization mechanisms for diverse GPCRs complements the GPCR signaling partner couplings profiles, including several types of G proteins and βarrs, in GPCRdb (https://gpcrdb.org/) ^58, 59^.

The extensive dataset shown here provides a valuable resource for research in GPCR signaling. Although internalization was initially linked to receptor desensitization and resensitization, compelling pieces of evidence indicate that the receptors can induce various signaling within cellular compartments^60–63^. It has been demonstrated that some GPCRs mediate sustained G protein signaling^64^ and activate the βarrs-dependent mitogen-activated protein kinases at endosome^14^. Recently, the activation of β_2_AR was shown to induce extracellular-signal-regulated kinase (ERK) activity originating from endosomes, but not the plasma membrane, and progress to the nucleus to regulate transcription^65^. Knowing the internalization profile for individual GPCR, which is highlighted in this dataset, is useful for controlling GPCR downstream signaling, spatially and temporally.

Classically, βarrs are believed to be the main regulators for GPCR internalization. Revising this view, we found βarrs are dispensable for GPCR constitutive internalization and are only required for 32% of these 60 receptors’ agonist-induced internalization (Supplementary figure and Figure). Prior to this work, the individual βarr contribution to GPCR internalization was debated^37–41^. Due to the lower expression level of βarr1 in the most used HEK293 cells, its contribution to GPCR internalization is largely undervalued (Figure 1). By overexpression of each βarr in Δβarr1/2 cell line, both βarrs show a similar overview effect on receptor internalization, although βarr2 has stronger effects in some cases^45^ (Figure 2). Indeed, the expression patterns can affect βarrs function. In Hep G2 cells, the endogenous βarr1 can drive several receptor internalizations (Figure 1). Similarly, βarr1 facilitates cancer cell, MDA-MB231, survival during hypoxia^36^.

The presence of multiple pathways for internalization provides the cell with flexibility and robustness. It ensures that even if one pathway is blocked, the cellular process can still occur through alternative routes. Degeneracy in the biological system is a ubiquitous property at every level of biological organization, from the genetic code level up to the interanimal communication level^42^. In the cellular system, various pathways might evolve to regulate one specific receptor internalization. For more than 30% of these 60 GPCRs, βarrs are partially required for their agonist-induced internalization (Figure 2), indicating other mechanisms are also involved ^17^. GLP-1R, as well as the protease-activated receptor 1 (PAR1), are examples that serve to show the salience of the degeneracy concept (Figure 7G), which were reported to recruit βarrs but do not need βarrs for agonist-induced internalization^6, 8, 46^. Indeed, βarrs take over for GLP-1R internalization, once the main process, the βarrs-independent way, was inhibited by 6A mutations (Figure 5D) or the natural genetic variants (Figure 8). More interestingly, although GRKs are required for βarrs-dependent and independent internalization processes, a GRK-independent way is also observed for GLP-1R internalization (Figure 7), which might be regulated by caveolin-1^54^.

This work also illustrates that the clathrin-mediated path plays a predominant role in GLP-1R internalization, which is the best-characterized endocytic route in mammalian cells^33,66^. GLP-1R can recruit clathrin adaptor AP2 in both βarrs-dependent and independent manners for agonist-induced internalization (Figure 7G). In agreement with the canonical βarr-dependent, clathrin-mediated internalization process, upon agonist stimulation, the βarr phosphorylation bar code, located at the distal CTD of GLP-1R, is phosphorylated by GRKs for βarr recruitment (Figure 6 and 7). After recruiting, the autoinhibitory C-terminus of βarr is displaced by phosphorylated receptor CTD, exposing an accessible site (DxxFxxFxxxR motif) to interact with the β2-adaptin subunit of the AP2 complex^44, 67–70^.

Alternatively, the μ2-adaptin subunit of AP2 has been shown to interact directly with several GPCRs through different motifs^19, 46, 71^, including the canonical tyrosine cargo internalization motif (YXXØ, Ø hydrophobic residue)^72, 73^ present at the extreme CTD of PAR1^46^ and an unusual stretch of eight arginine residues (rather than the YXXØ) within the CTD of the α_1b_-adrenergic receptor^71^. However, in the case of GLP-1R, we reveal a novel motif, TSSLSS, within the middle CTD. This region probably binds to μ2 subunits of AP2^32^, which need to be further studied. The downregulation of μ2 subunits substantially inhibits agonist-induce GLP-1R internalization (Figure). Furthermore, the TSSLSS region might undergo post-translation modification by GRKs for the βarr-independent AP2 recruitment (Figure 7). More studies are needed to check whether this region is phosphorylated or not, as GRKs can function without phosphorylating the receptor^74, 75^. Together, these findings suggest that the clathrin adaptor AP2 functions in distinct modes for GPCR internalization. To our surprise, distinct modes, both βarrs-dependent and independent manners, cooperate in one receptor, GLP-1R. The precise mechanism for GLP-1R internalization machinery, including GLP-1R/AP2 complex, GLP-1R/βarr complex, and GLP-1R/βarr/AP2 mega-complex, need to be further clarified by structural studies.

Several mutations in the AP2 complexes have been identified in patients with a range of genetic disorders^76^. Missense mutations of the AP2 σ subunit impair a G-protein coupled calcium-sensing receptor (CaSR) endocytosis, resulting in familial hypocalciuric hypercalcemia type 3^77, 78^. A recurrent missense mutation in AP2 μ2 subunits impairs clathrin-mediated endocytosis and causes developmental and epileptic encephalopathy (DEE)^9^. Notably, a DEE patient was found to carry a homozygous mutation in the *GLP1R* gene^79^. These suggest that targeting the GLP-1R clathrin-mediated internalization process might be a treatment for DEE. Indeed, GLP-1 analogs have been reported to exert neuroprotective effects in mouse models of acute and chronic epilepsy^80–82^. In addition, GLP-1R internalization is essential for regulating insulin secretion and blood glucose homeostasis^31, 32^. Interestingly, several natural genetic variants at the CTD of GLP-1R, which impair the βarr-independent clathrin-mediated internalization (Figure 8), were not reported to link to any diseases, until now^56^. For the people who carry these variants, βarrs might take over for GLP-1R internalization (Figure 8) to keep them healthy, supporting the concept of degeneracy in the biological system.

In conclusion, we systematically reexamine the contribution of βarrs to an unprecedented number of GPCR internalization and provide insight into the mechanism for GLP-1R internalization, which may also apply to other GPCRs in general. Our studies revise the canonical concept of the GPCR internalization process, and open an opportune pharmacological intervention window for controlling various signaling events, providing considerable possibilities for new therapies.

## Materials and Methods

### Materials

Agonists for the 60 GPCRs were purchased from Tocris Bioscience (Ellisville, MO, USA), or Sigma-Aldrich (St. Louis, MO, USA). Lipofectamine 2000 (11668019) were obtained from Life Technologies (Carlsbad, CA, USA). Fluorescein sodium (46955), were purchased from Sigma-Aldrich (St. Louis, MO, USA). SNAP-Lumi4-Tb (SHALOTBC) labeling reagents, total β-arrestin1 (64BAR1TPEB), total β-arrestin2 (64BAR2TPEB), and β-arrestin2/AP2 interaction (62BDBAR2PEB) cellular kits were from Revvity Cisbio (Codolet, France). Barbadin (HY-119706) was purchased from MedChemExpress (NJ, USA).

### Plasmids

The pcDNA3.1 plasmids encoding wild-type 60 human GPCRs, respectively, tagged with SNAP, were from Revvity Cisbio. The pRK5 plasmids encoding wild-type human GLP-1R, M2R, MOR, V1AR, V2R, and β_2_AR were tagged with a double tag, HA-SNAP, inserted immediately after the signal peptide^83^. Site-directed mutations in the plasmids were generated using the QuikChange mutagenesis protocol (Agilent Technologies). The probes (full length of EGFP) were fused to the CTD of GLP-1R and MOR, respectively, with AflII restriction site as a linker^84^. The plasmids encoding human βarr1, βarr2, GRK2, GRK3, GRK5, and GRK6 in pcDNA3.1 were tagged with Flag-TST (provided by ARPEGE platform, IGF). cDNA encoding chimeric human GLP-1R_MO_, GLP-1R_M2_, GLP-1R_V1A_, GLP-1R_V2_, GLP-1R_β2A_, MOR_GLP_, M2R_GLP_, V1AR_GLP_, V2R_GLP_, and β_2_AR_GLP_ were synthesized directly and subcloned into pRK5 plasmids fused with a double tag, HA-SNAP. The sequence coding GLP-1R CTD (RDSSMKPLKCPTSSLSSGATAGSSMYTATCQASCS) were exchanged with the CTDs from MOR (IPTSSNIEQQNSTRIRQNTRDHPSTANTVDRTNHQLENLEAETAPLP), M2R (CHYKNIGATR), V1AR (MKEKFNKEDTDSMSRRQTFYSNNRSPTNSTGMWKDSPKSSKSIKFIPVST), V2R (GRTPPSLGPQDESCTTASSSLAKDTSS), β_2_AR (RRSSLKAYGNGYSSNGNTGEQSGYHVEQEKENKLLCEDLPGTEDFVGHQGTVPSDNIDSQGRNCSTNDSLL), respectively, to make these chimeric receptors.

### Cell culture and transfection

Hep G2 cells (ATCC, HB-8065), HEK293 cells (ATCC, CRL-1573), GRK2/3/5/6 knockout HEK293 cells^45^ (ΔGRK2/3/5/6 cells, were kind gifts from Dr. Carsten Hoffmann (Universitätsklinikum Jena, Jena, Germany)), and βarr1/2 knockout HEK293 cells^38^ (Δβarr1/2 cells, were kind gifts from Dr. Asuka Inoue (Tohoku University, Sendai, Miyagi, Japan)) were cultured in Dulbecco’s modified Eagle’s medium (DMEM) supplemented with 10 % FBS and 100 U/ml Penicillin–Streptomycin at 37°C and 5 % CO_2_ in a humidified incubator. Plasmids were transfected by Lipofectamine 2000 as manufacturer’s recommendations. On-TARGETplus Smartpool siRNAs target βarr1 (L-011971-00-0020), βarr2 (L-007292-00-0020), AP2 μ2 (L-008170-00-0005), caveolin-1 (L-003467-00-0005), caveolin-2 (L-010958-00-0005), caveolin-3 (L-011229-00-0005), and ARF6 (L-004008-00-0005), respectively, were from Horizon. A Non-targeting pool (D-001810-10-20) was used as control. 60-70 % confluent cells were transfected with both siRNAs (100 nM) and plasmids (100 ng/well, 96-wells plate) using DharmaFECT1 transfection reagent (T-2001-03) as manufacturer’s recommendations. Data are collected in Hep G2 cells, HEK293 cells, ΔGRK2/3/5/6 cells or Δβarr1/2 cells as stated in figure legends. Cells were regularly checked for mycoplasma infections using the LONZA MycoAlert mycoplasma detection kit (LT07-318).

### β-arrestin1 and β-arrestin2 expression

βarr1 and βarr2 expression in cells was measured using a total βarr1 and βarr2 cellular kit (Revvity Cisbio), according to the manufacturer’s recommendations. TR-FRET measurements were performed using a PHERAstar FS microplate reader (BMG Labtech, Ortenberg, Germany). After excitation with a laser at 337 nm (40 flashes per well), the fluorescence was collected at 620 nm and 665 nm for a 400 μs reading after a 60 μs delay after excitation. The ratio between 665 nm and 620 nm was determined and then plotting it.

### Diffusion-enhanced resonance energy transfer internalization assay

A diffusion-enhanced resonance energy transfer (DERET) internalization assay was performed for measuring GPCR internalization in real time^43^. Transfected cells in black non-transparent 96-well plates were labeled in Tag-lite labeling buffer (Revvity Cisbio) with 100 nM SNAP-Lumi4-Tb for 1 h at 4°C. Excess of Lumi4-Tb was removed by washing each well three times with 100 μl of Tag-lite labeling buffer. Internalization experiments were performed by incubating the cells with Tag-lite labeling medium, either alone or containing agonist for the receptor, in the presence of fluorescein. Typically, in plates containing Lumi4-Tb-labeled cells, 50 μl of buffer containing agonist at the indicated concentrations was added, immediately followed by the addition of 50 μl of 48 μM fluorescein. Afterwards, the internalization kinetics were recorded at 37°C.

Lumi4-Tb was excited by a laser at 337 nm, and the emission fluorescence intensities were recorded for the donor (620 nm, 1500 μs delay, 1500 μs reading time) and acceptor (520 nm, 150 μs delay, 400 μs reading time) using a PHERAstar FS microplate reader. The ratio of 620/520 was obtained by dividing the donor signal (620 nm) by the acceptor signal (520 nm) and multiplying this value by 10,000.

### Fluorescence labeling for quantifying cell surface receptor expression

For cell surface receptor expression quantification, SNAP-tagged GPCRs were labeled with 100 nM SNAP-Lumi4-Tb at 4°C for 1 h. After being labeled, cells were washed twice with Tag-lite buffer. Afterwards, the cell surface receptor expression was recorded. Lumi4-Tb was excited by a laser at 337 nm, and the emission fluorescence intensities were recorded for the donor (620 nm, 60 μs delay, 400 μs reading time) using a PHERAstar FS microplate reader.

### GLP-1R internalization in mouse pancreatic β-cells

As already reported^85^, the heterozygous (*Arrb2^+/−^*) mice were obtained from R. J. Lefkowitz (Duke University Medical Center, Durham, NC, USA). Mice were subsequently backcrossed onto C57BL/6J background (Charles River, Lyon, France). *Arrb2^−/−^* mice and their wild-type littermates (*Arrb2^+/+^*) were generated by breeding heterozygous animals, and genotyping was performed by PCR^85^. Studies complied with the animal welfare guidelines of the European Community (EU directive 2010/63/EU) and was approved by the ethic committee (CEEA N°36) and the ministry of education and research, France (D34-172-41). Mice were housed in pathogen free animal house, with a 12light/12dark cycle, with a temperature of 20°C, and had freely access to food (4% Fat diet) and water. All experiments were performed using 4-5 month-old male mice. Islets were isolated after collagenase digestion of the pancreas^85, 86^, dispersed in clusters of cells with trypsin and plated on glass coverslips in RPMI-1640. For the experiments, the medium was a bicarbonate buffered (KRB) solution containing (mM): NaCl 120, KCl 4.8, CaCl2 2.5, MgCl2 1.2 and NaHCO3 24. It was gassed with O2:CO2 (95:5) to maintain pH 7.4 and supplemented with 1 mg/ml BSA. Afet two days culture, dispersed mouse islet cells were preincubated for 15min in 8mM glucose (G8) before being incubated for 25min in the presence or absence of agonists (GLP-1 or Ex-4), as indicated. Cells were fixed for 20min (4% paraformaldehyde), overnight incubated with primary antibodies (MAB7F38; 5µg/ml; DSHB) followed by 1h incubation with secondary antibodies (#715-166-150, Cy3-conjugated AffiniPure, Donkey anti-mouse, dilution 1/4000, Jackson Immuno Research) and DAPI. Cells were imaged using an x100 objective from AxioImager Apotome microscope (Zeiss) using the ZenBlue software. All of the images from *Arrb2^+/+^* and *Arrb2^-/-^* and different conditions (in the presence or absence of agonists) from the same islet preparation were taken the same day. 30-50 cells per coverslip/mouse (n=1) were imaged and saved in “.czi” (rather than “tiff”) format allowing to save the entire image parameters and were quantified using ImageJ, as already reported^86^. Each dot represent the average of ∼30-50 cells from 1 mouse. Cells were pre-incubated 15min with barbadin (30µM) prior the experiments.

### βarr2 and AP2 recruitment

A TR-FRET assay was performed for measuring βarr2 or AP2 recruitment by receptors. EGFP-fused receptors without or with βarr2, as stated in figure legends, were transfected in white non-transparent 96-well plates for 24 h. After stimulating the cells with agonist at the indicated concentrations at room temperature for 20 min, the cells were fixed with stabilization buffer (Revvity Cisbio) for 15 min. After three times washing with wash buffer (Revvity Cisbio), cells were incubated with the anti-βarr2-Tb antibody (Revvity Cisbio) or anti-AP2-Tb antibody (Revvity Cisbio) at final concentration of 0.5 nM for 2 h. Afterwards, the HTRF ratio were recorded using a PHERAstar FS microplate reader.

Lumi4-Tb was excited by a laser at 337 nm, and the emission fluorescence intensities were recorded for the donor (620 nm, 60 μs delay, 400 μs reading time) and acceptor (520 nm, 60 μs delay, 400 μs reading time). The ratio of 520/620 was obtained by dividing the acceptor signal (520 nm) by the donor signal (620 nm) and multiplying this value by 10,000.

### βarr2/AP2 interaction assay

βarr2/AP2 interaction in cells was measured using a β-arrestin2/AP2 interaction (62BDBAR2PEB) cellular kit (Revvity Cisbio), according to the manufacturer’s recommendations. TR-FRET measurements were performed using a PHERAstar FS microplate reader. After excitation with a laser at 337 nm, the fluorescence was collected at 620 nm and 665 nm for a 400 μs reading after a 60 μs delay after excitation. The ratio between 665 nm and 620 nm was determined and then plotting it.

### Western blot

Cells were washed once with ice-cold PBS and subsequently lysed with RIPA Buffer (ThermoFisher, 89900), supplemented with protease inhibitor cocktails (Merck, 5892953001). Cleared lysates were boiled with sodium dodecyl sulfate (SDS) loading buffer and 15 μg of total protein were loaded onto each lane of 8% polyacrylamide gels. After transfer onto nitrocellulose membranes, the total protein was detected by using specific antibodies (AP2: mouse anti-AP2 μ2, BD Biosciences, 611351, (1:250), Actin: mouse anti-β actin, Sigma-Aldrich, A5441(1:5000)). As secondary antibodies, we used rabbit anti mouse IgG-d2, Revvity Cisbio, 61PAMDAB (1:100).

### Curve fitting and data analysis

All data in figures and supplementary figures are mean ± SEM of at least three independent experiments performed in triplicate, unless stated differently in the figure legends. The curves were fitted using Prism software (Version 9, GraphPad Software). Fluorescent images were analyzed using lmageJ (Version 1.440, National Institutes of Health). Statistical differences were calculated using GraphPad Prism. P-values were determined using unpaired two-tailed t-test, paired two-tailed t-test, or one-way ANOVA with Dunnett’s multiple comparisons test. Differences were considered to be no significantly different (ns) when P > 0.05. P-values were determined using paired, unpaired t-test or one-way ANOVA with Dunnett’s multiple comparisons test. ***p ≤ 0.001, **p ≤ 0.01, *p ≤ 0.05 and ns, not significant.

## Data Availability

Data supporting the findings of this manuscript are available from the corresponding authors upon reasonable request.

## Acknowledgments

We thank Dr. Asuka Inoue (Tohoku University, Sendai, Miyagi, Japan) for the β-arrestin1/2 KO cell line, and Revvity Cisbio (Codolet, France) for the reagents. We appreciate the ARPEGE (Pharmacology Screening-Interactome, Institut de Génomique Fonctionnelle, Montpellier) facility, for their valuable assistance in measurements. P. R. and J.-P. P. were supported by the Centre National de la Recherche Scientifique (CNRS; PRC n°1403), the Institut National de la Santé et de la Recherche Médicale (INSERM ; International Research Program « Brain Signal »), the Program Hubert Curien (PHC) Cai Yuanpei from the Ministère Français des Affaires Etrangères, and by grants from the Agence Nationale de la Recherche (ANR-13-RPIB-0009-01), the FRM (FRM team: DEQ20170336747).

## Author contributions

**Competing interests**: Philippe Rondard and Jean-Philippe Pin are involved in a collaborative team between the CNRS and Revvity Cisbio. The remaining authors declare no competing interests.

