## Supplementary figures for "Multi-faceted roles of β-arrestins in G protein-coupled receptors endocytosis"

### Supp Figures

Supp Fig. 1 expression profile of βarrs across native tissues and cell types

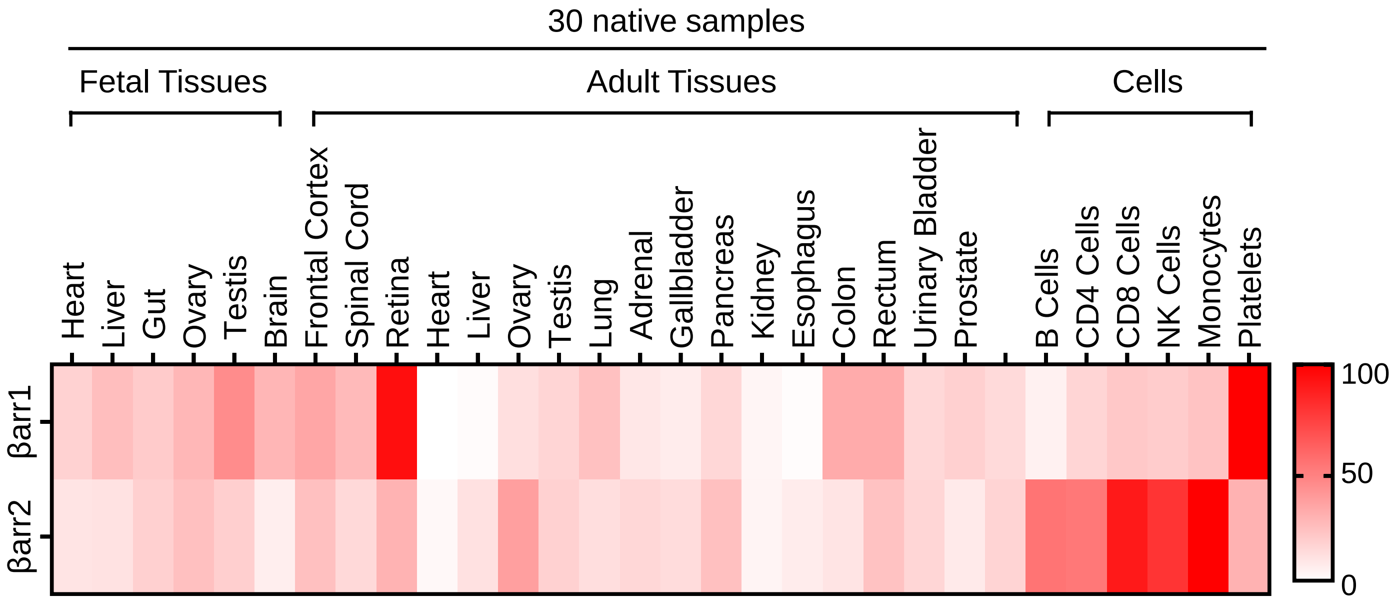

Supp Fig 2. Effect of βarrs siRNA on βarrs expression in HEK293 and Hep G2 cells

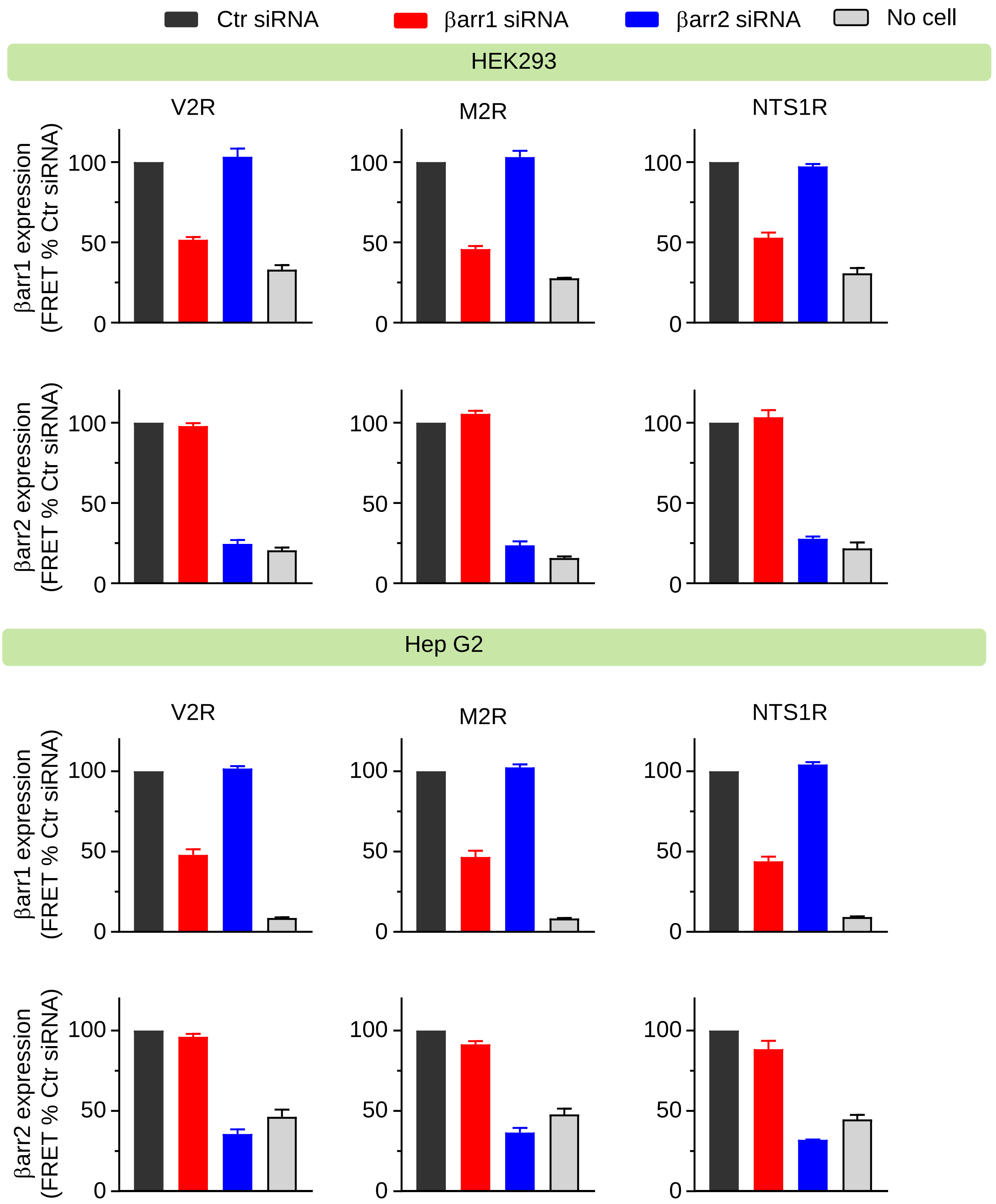

Supp Fig 3. Effect of βarrs siRNA on receptors cells surface expression in HEK293 and Hep G2 cells

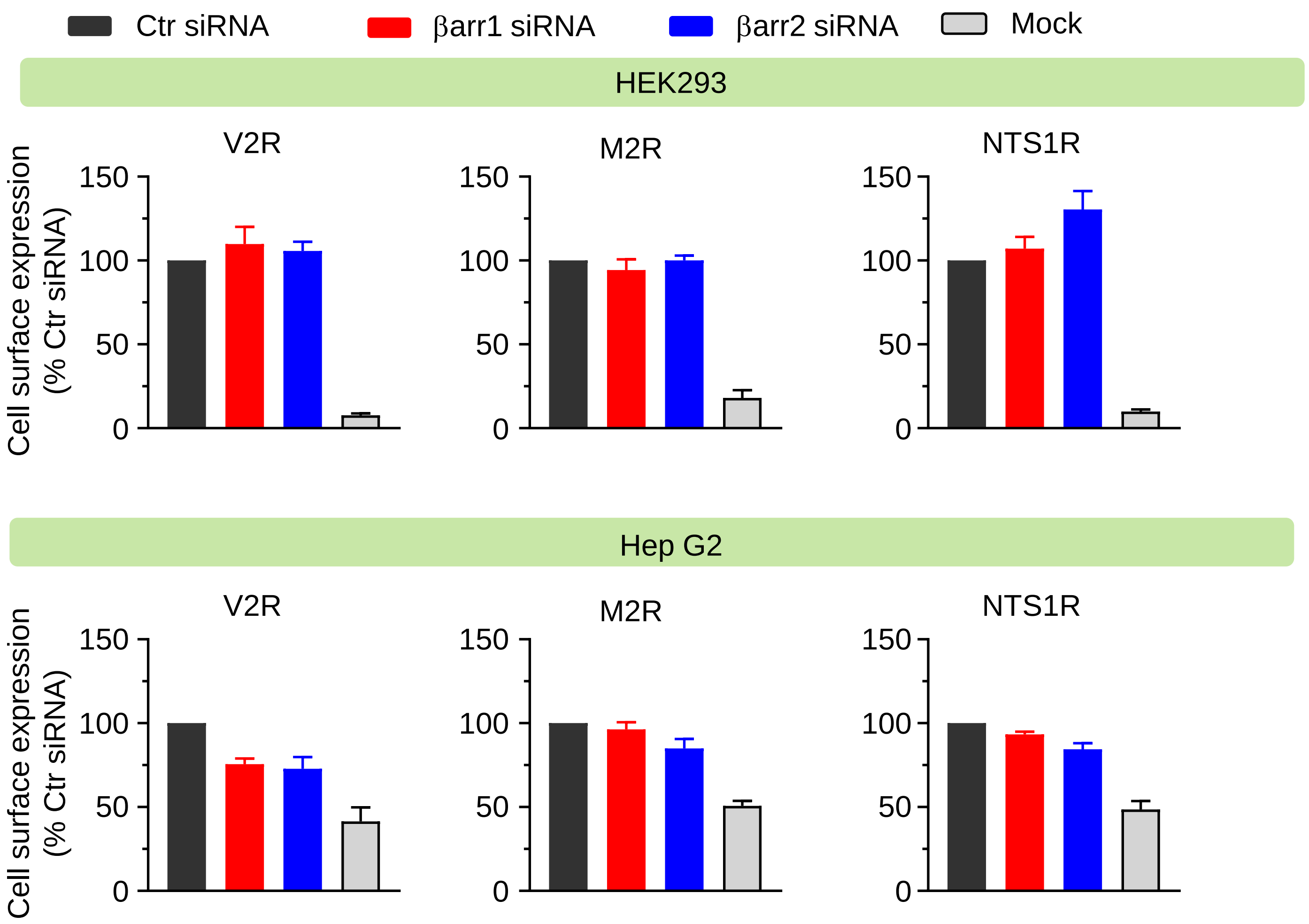

Supp Fig 4. Optimizing the transfection condition for βarr1 and βarr2.

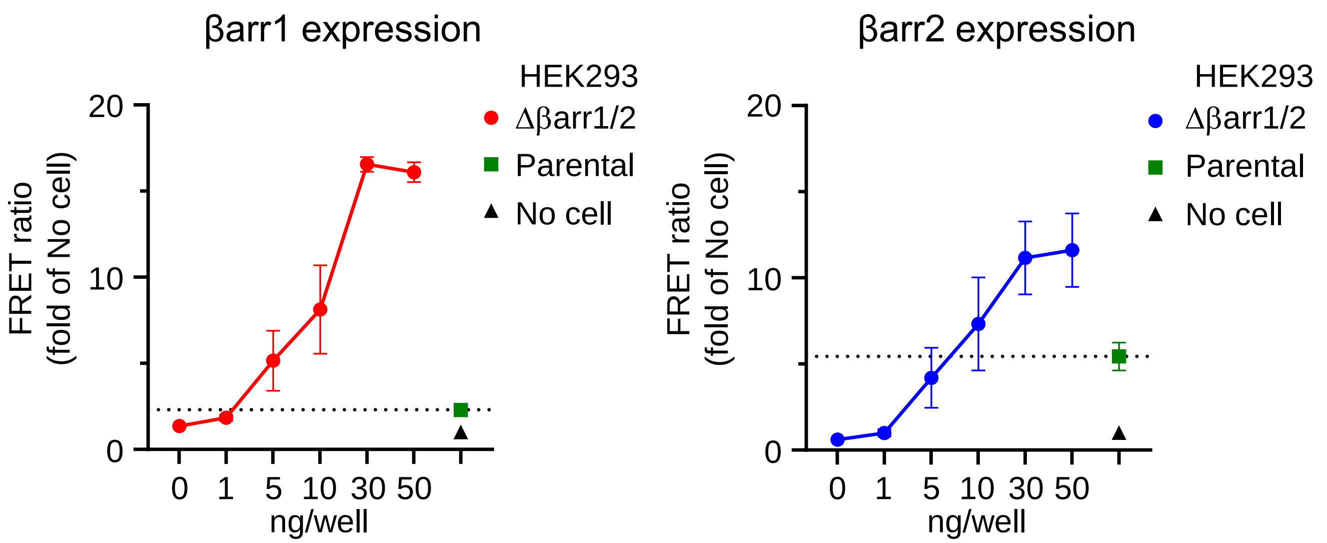

Supp Fig 5. Internalization profiles of 60 GPCRs

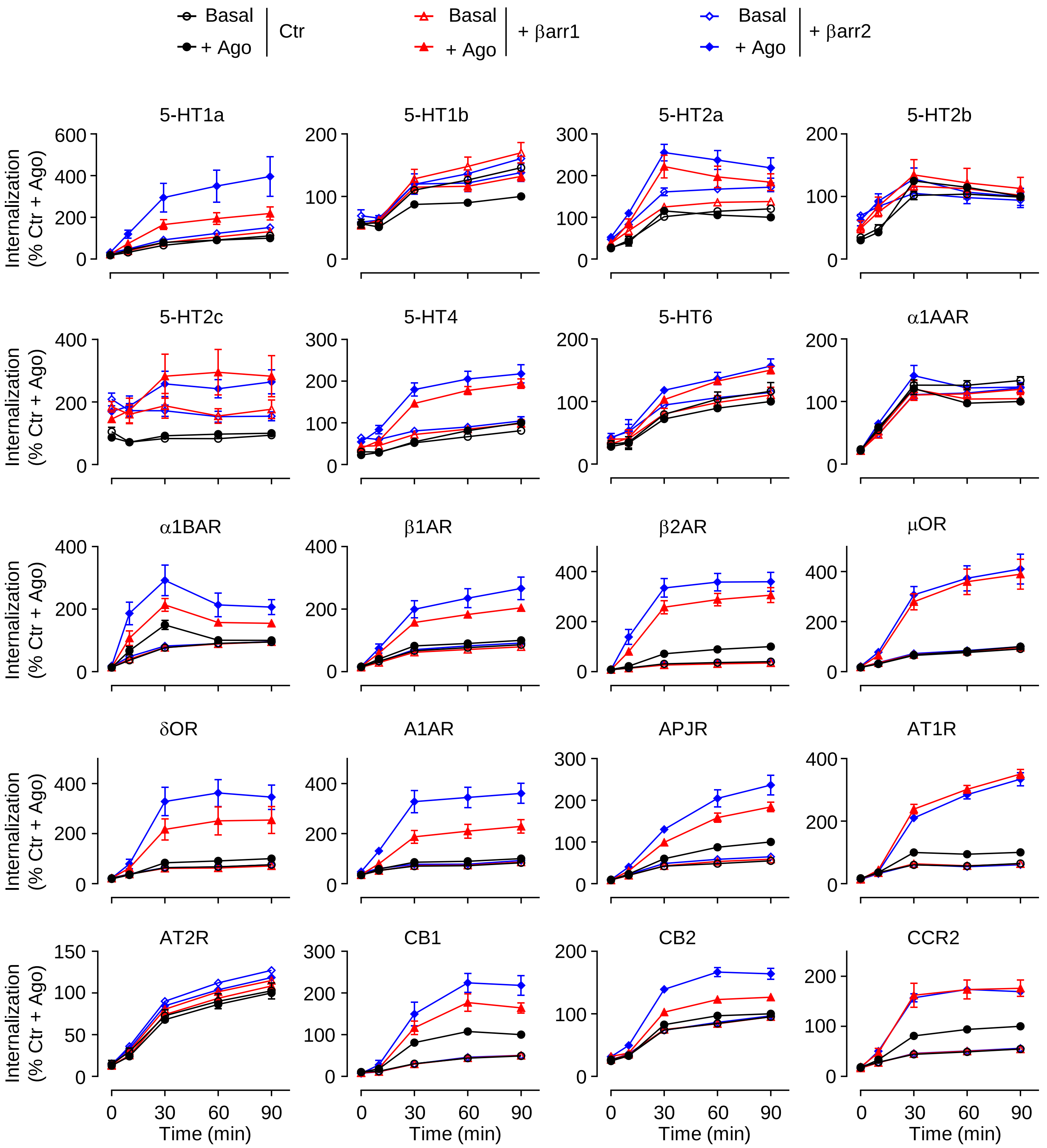

Supp Fig 5. Internalization profiles of 60 GPCRs

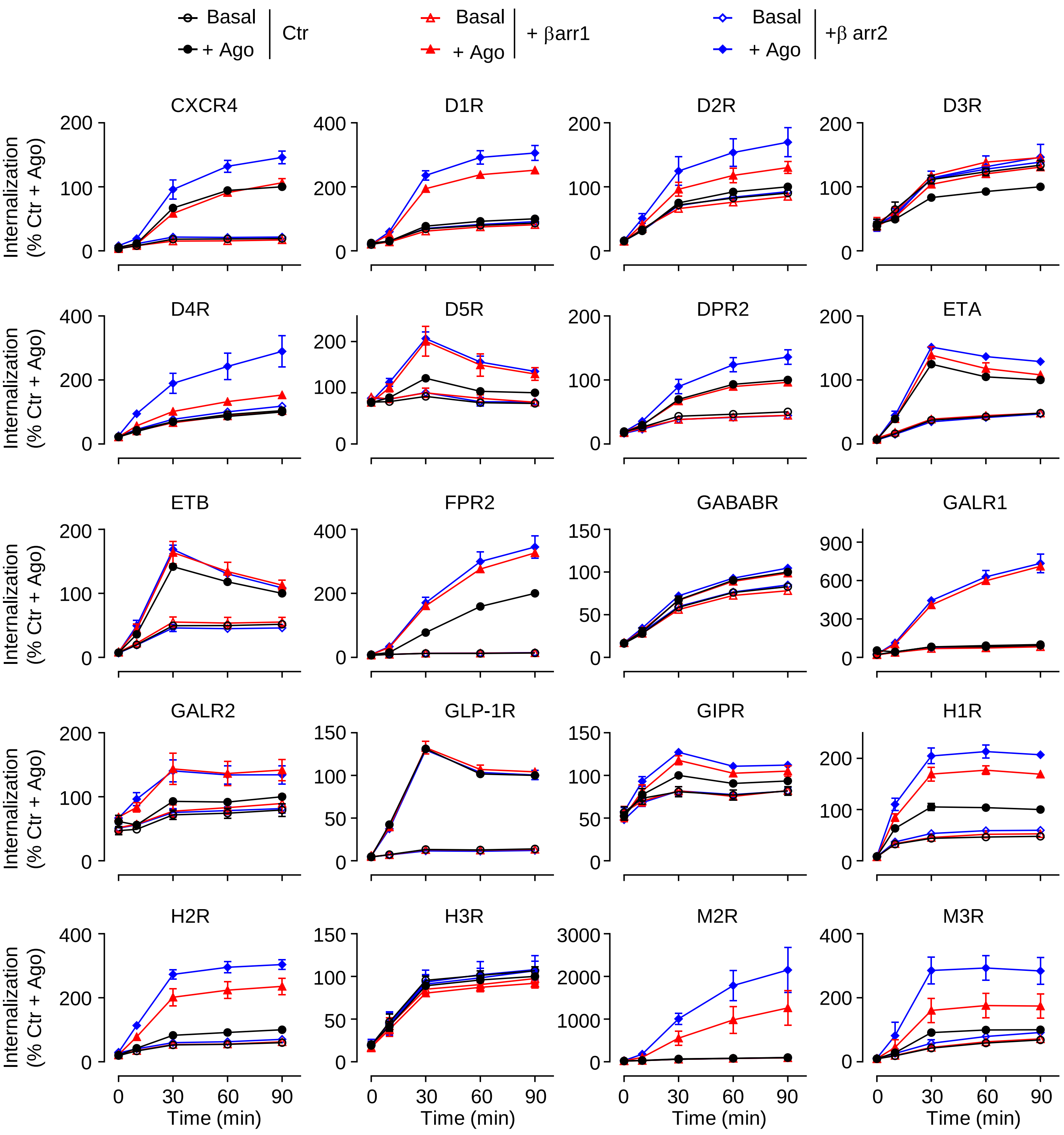

### Supp Fig 5. Internalization profiles of 60 GPCRs

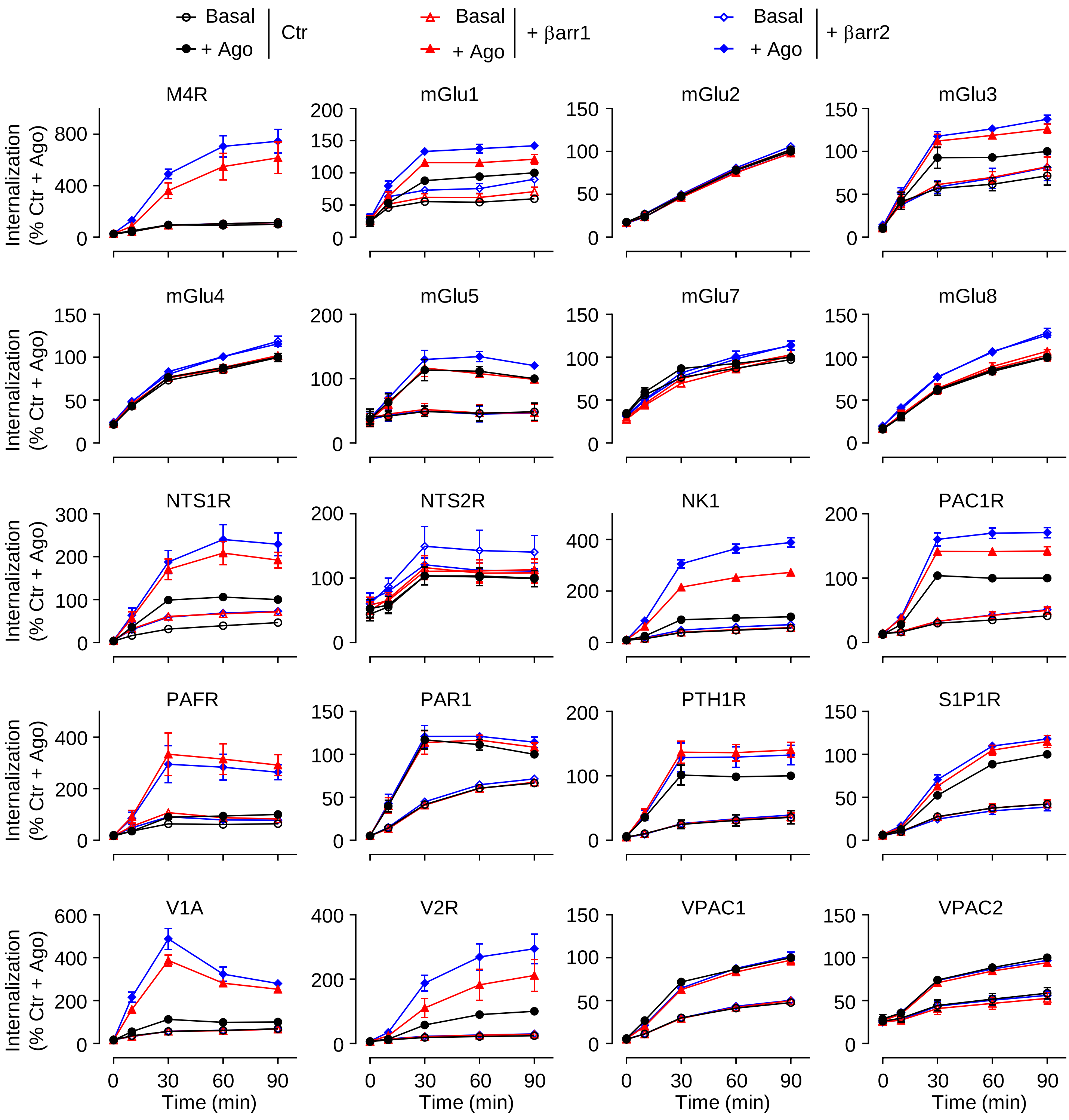

### Supp Fig 6. Constitutive internalization profiles of 60 GPCRs

#### Constitutive internalization (Δβarr1/2 cells)

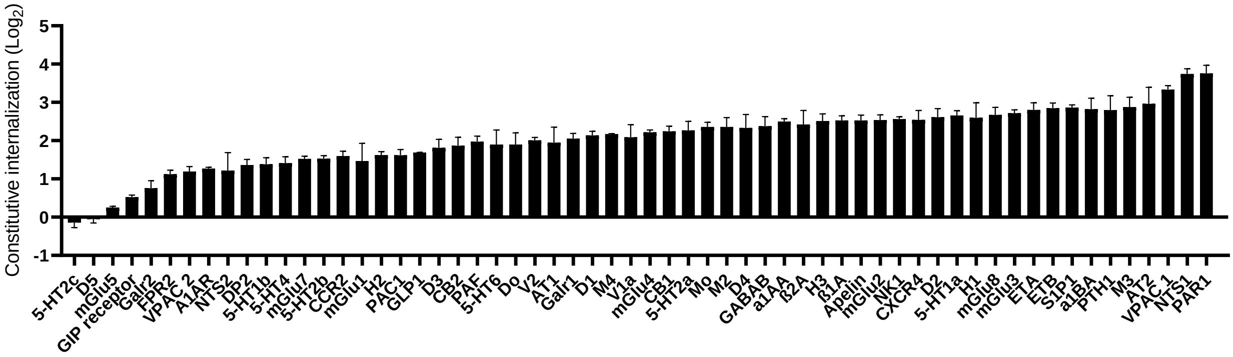


#### + βarr1

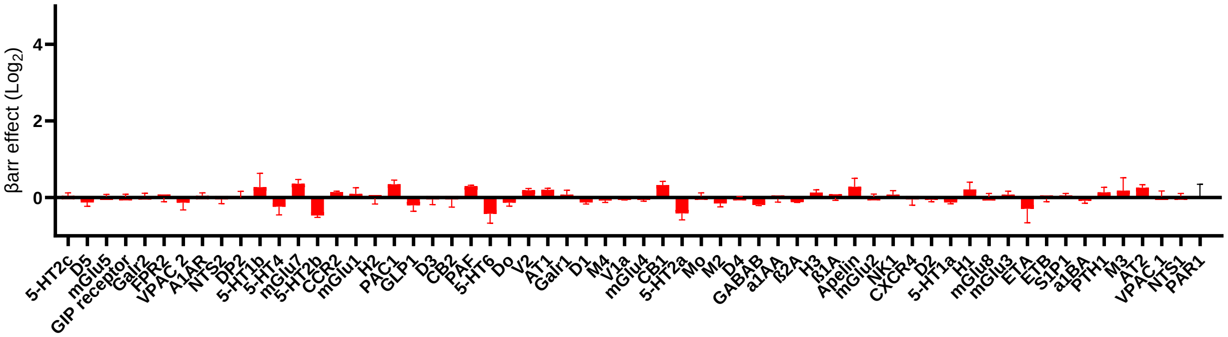


#### + βarr2

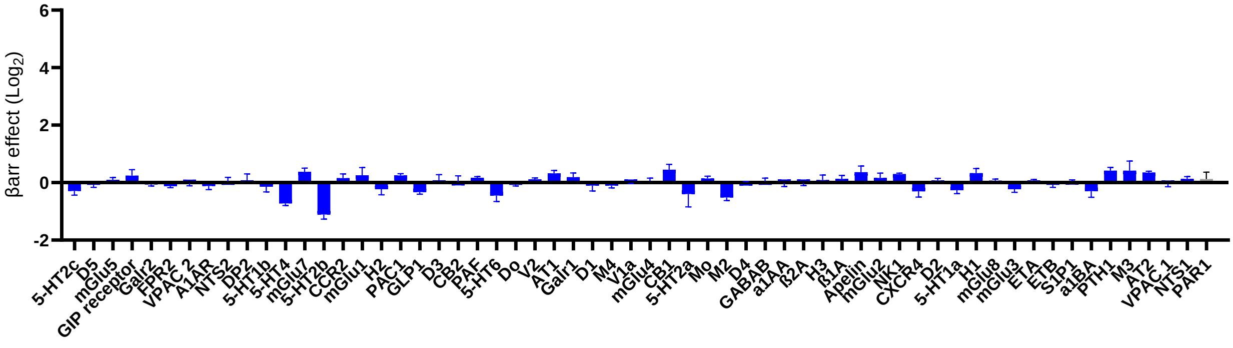

Supp Fig 7. GLP-1R and MOR expression in Δbarr1/2 HEK293 cells

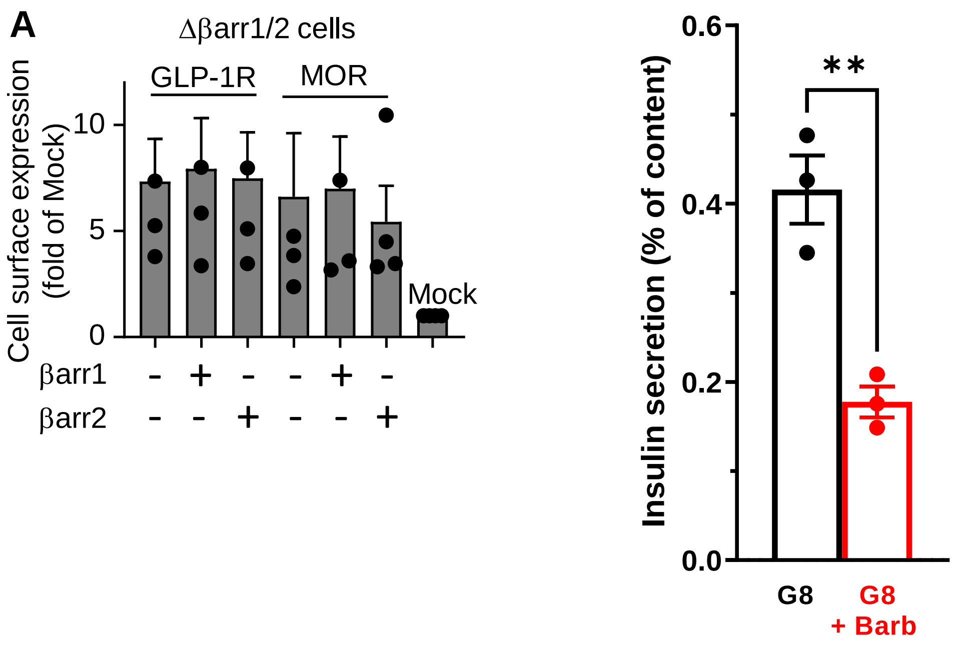

Supp Fig 8. Effect of barrs on GLP-1R and MOR internalization in HEK293 cells.

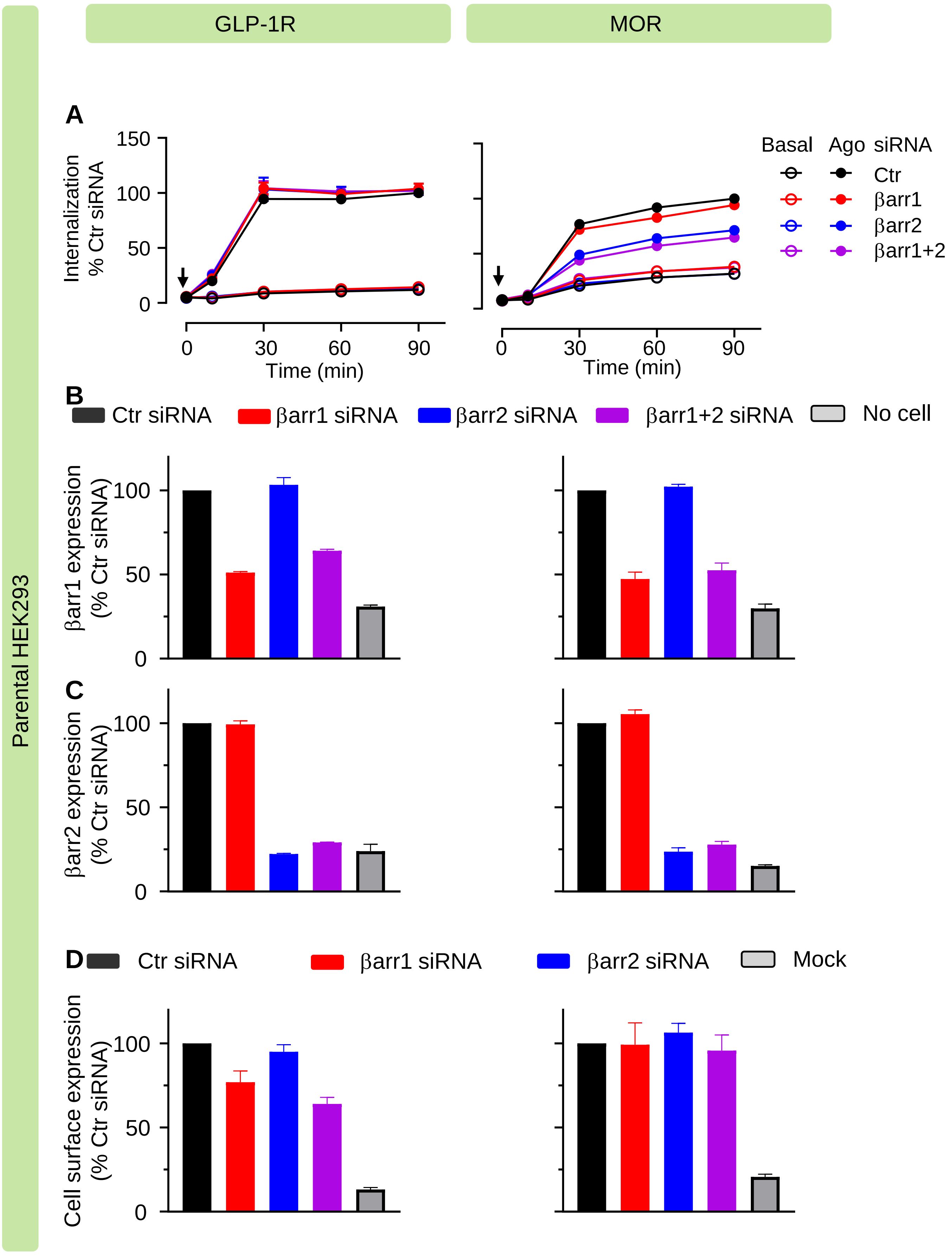

### Supp Fig 9. Effect of barrs on the internalization for the Chimeras

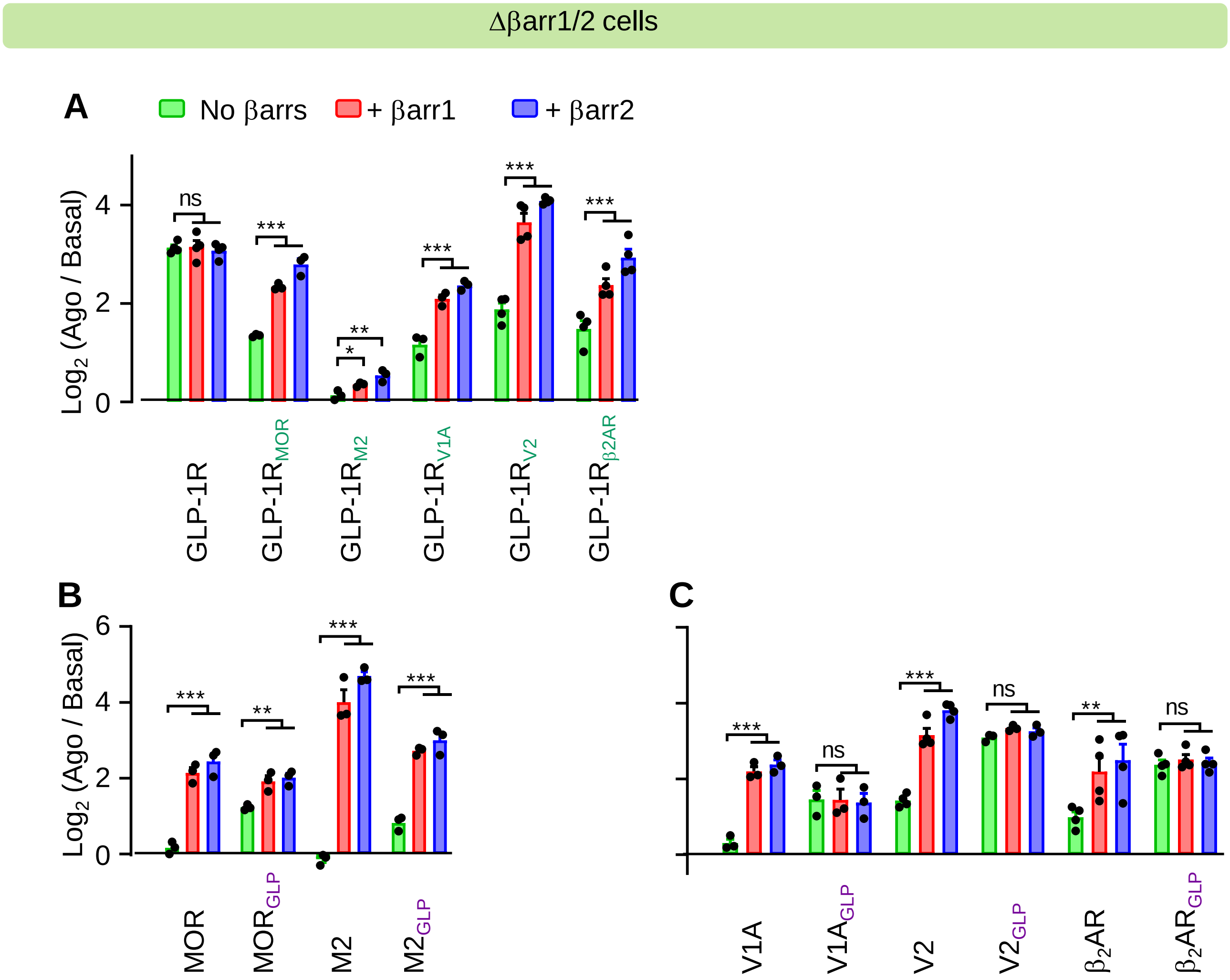

### Supp Fig 10. Internalization profiles of receptor chimeras

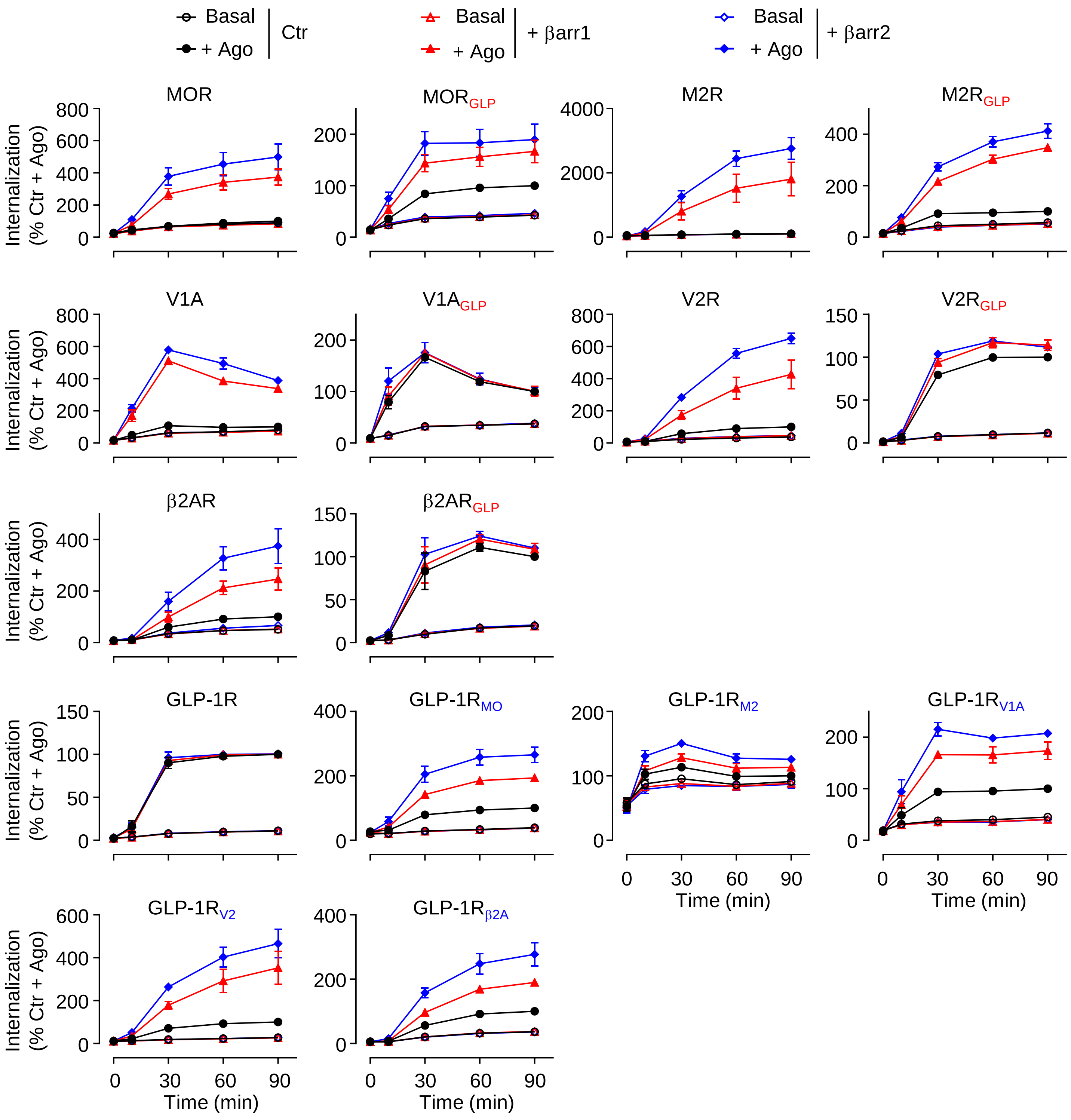

Supp Fig 11. Cell surface expression of receptor chimeras

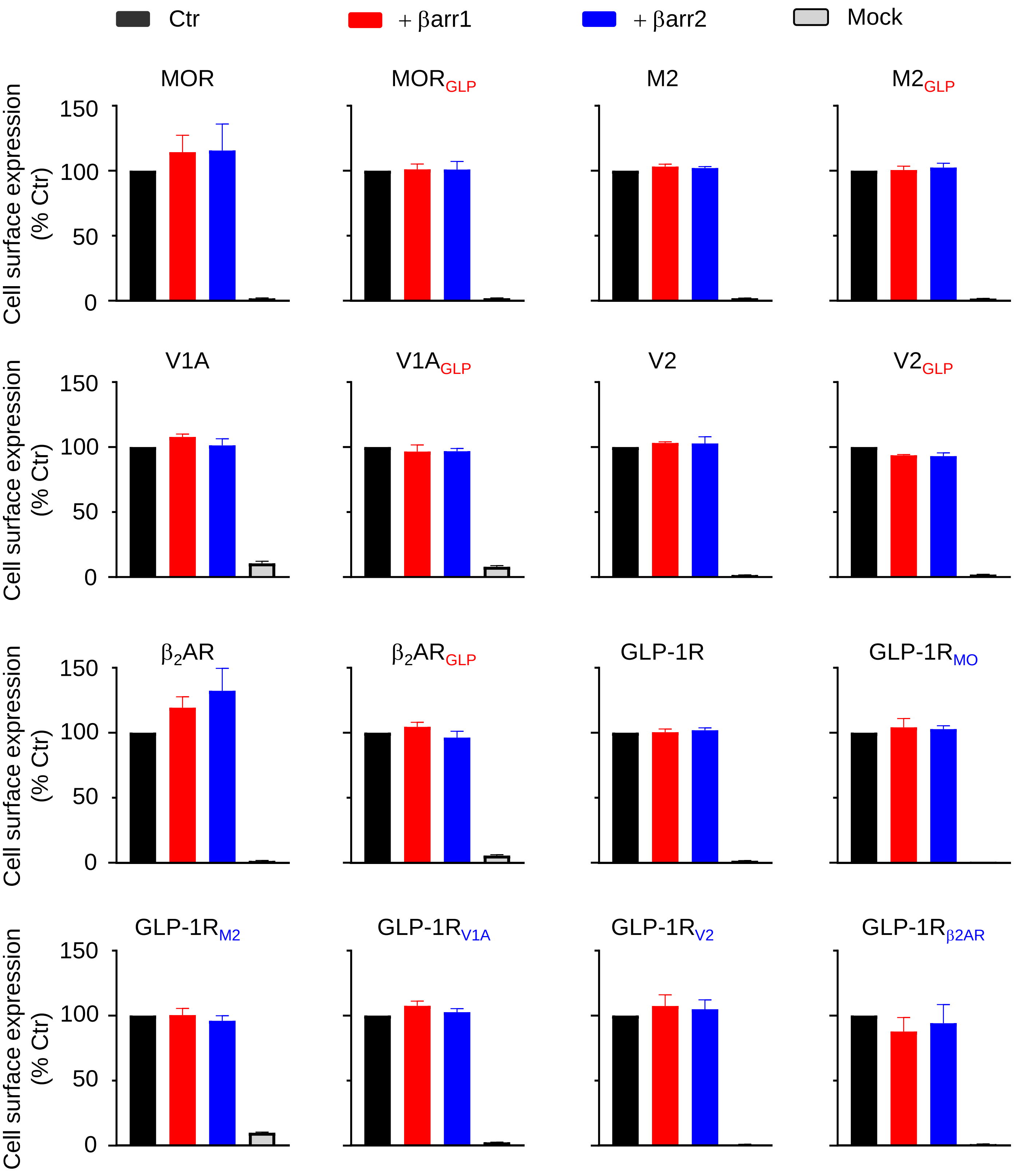

### Supp Fig 12. Amino acid alignment for CTD of GPCRs

| Receptors | C terminus sequence |
|---|---|
| GLP-1R | RDSSM KPLKC PTSSL SSGAT AGSSM YTATC QASCS |
| MOR | IPTSSN IEQQN STRIR QNTRD HPSTA NTVDR TNHQL ENLEA ETAPL P |
| M2 | CHYKN IGATR |
| V1A | MKEKF NKEDT DSMSR RQTFY SNNRS PTNST GMWKD SPKSS KSIKF IPVST |
| V2 | GRTPPS LGPQD ESCTT ASSSL AKDTS S |
| $\beta_2$AR | RRSSL KAYGN GYSSN GNTGE QSGYH VEQEK ENKLL CEDLP GTEDF VGHQG TVPSD NIDSQ GRNCS TNDSL L |

Supp Fig 13. Effect of siRNAs on GLP-1R internalization

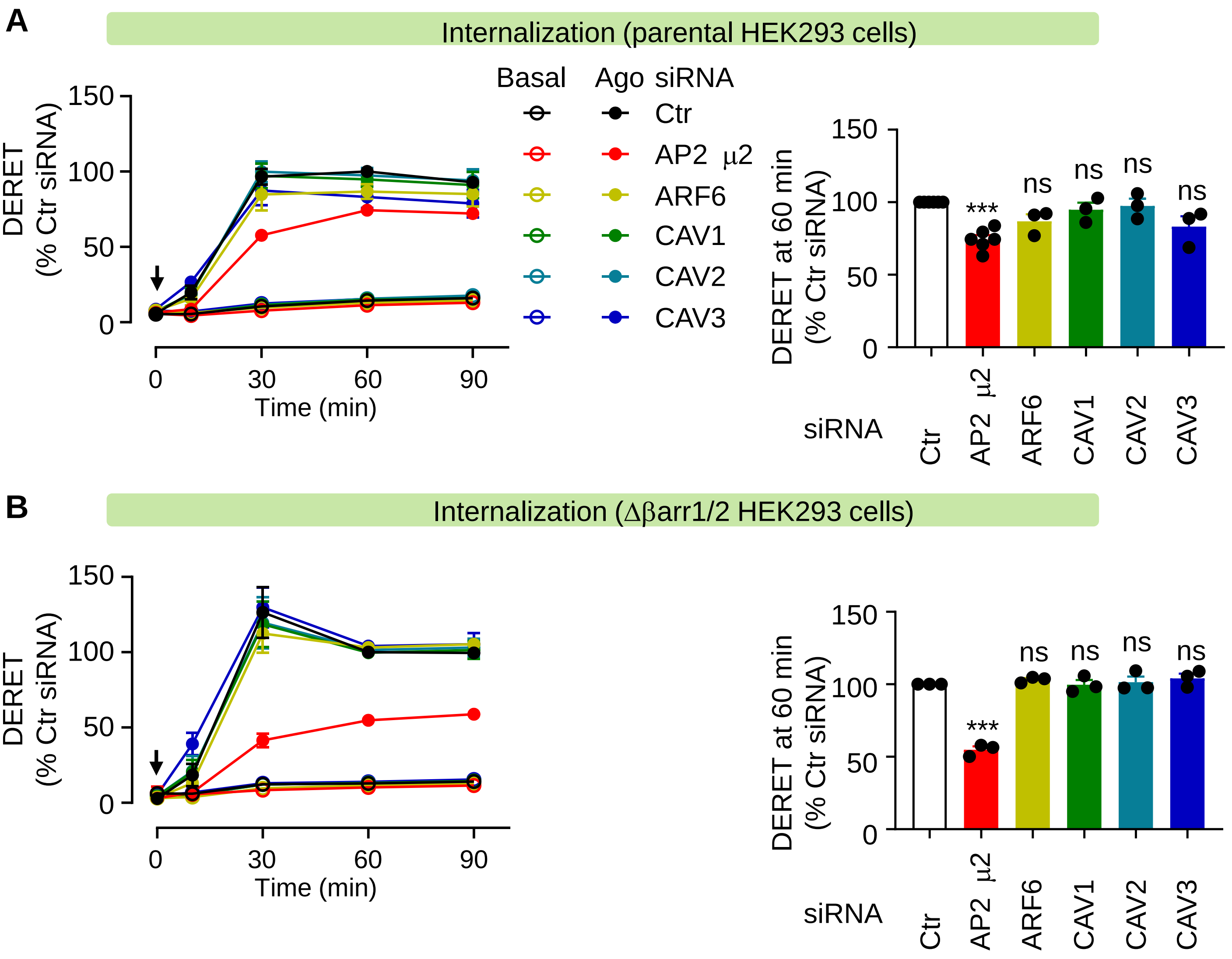

Supp Fig 14. Effect of combination of barrs and AP2 siRNAs on GLP-1R internalization

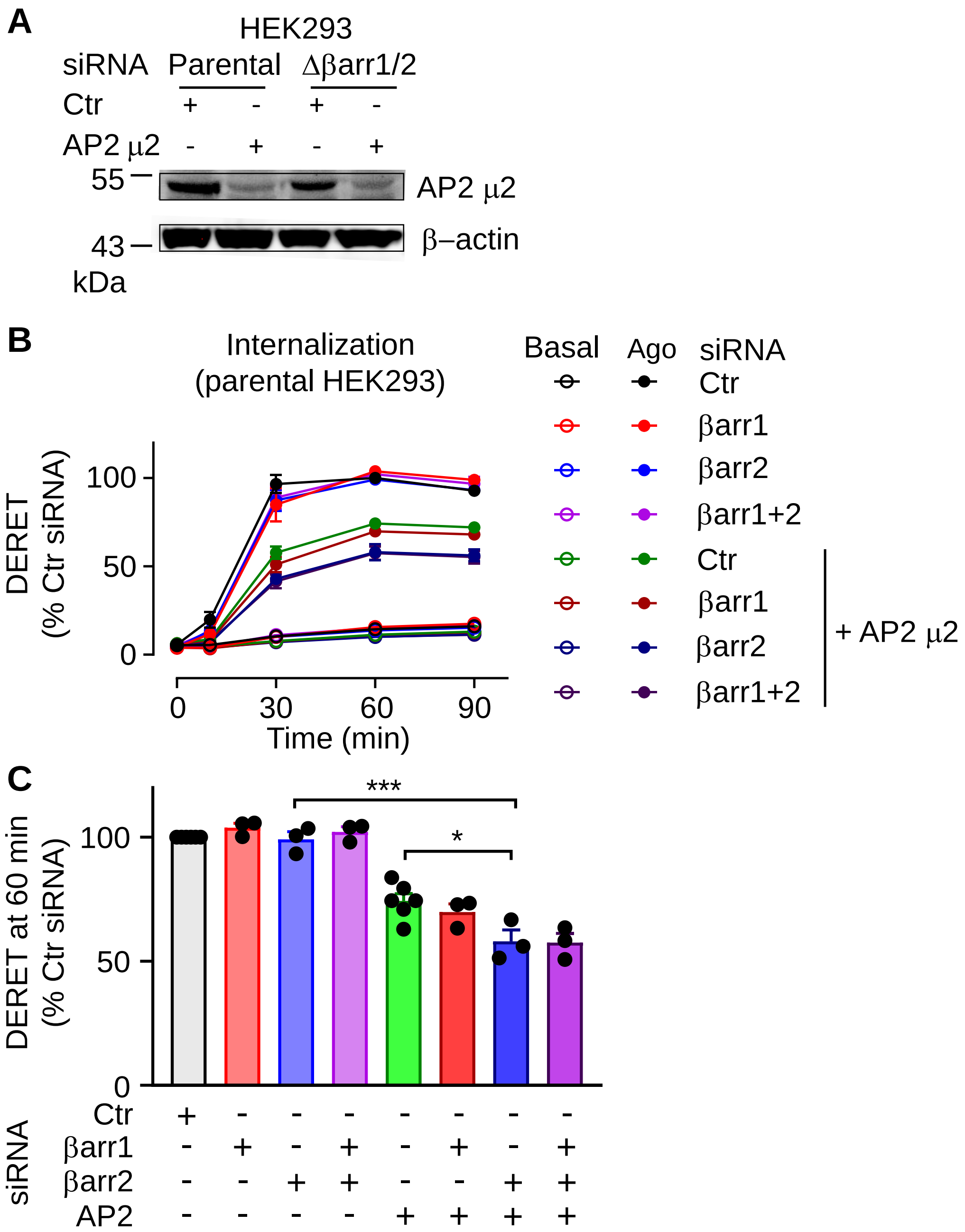

### Supp Fig 15. AP2 recruitment of MOR and GLP-1R.

AP2 recruitment (Δβarr1/2 HEK293 cells)

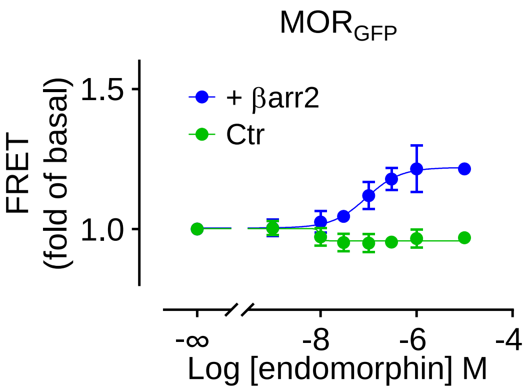

Supp Fig 16. βarrs effect on the internalization of GLP-1R mutants

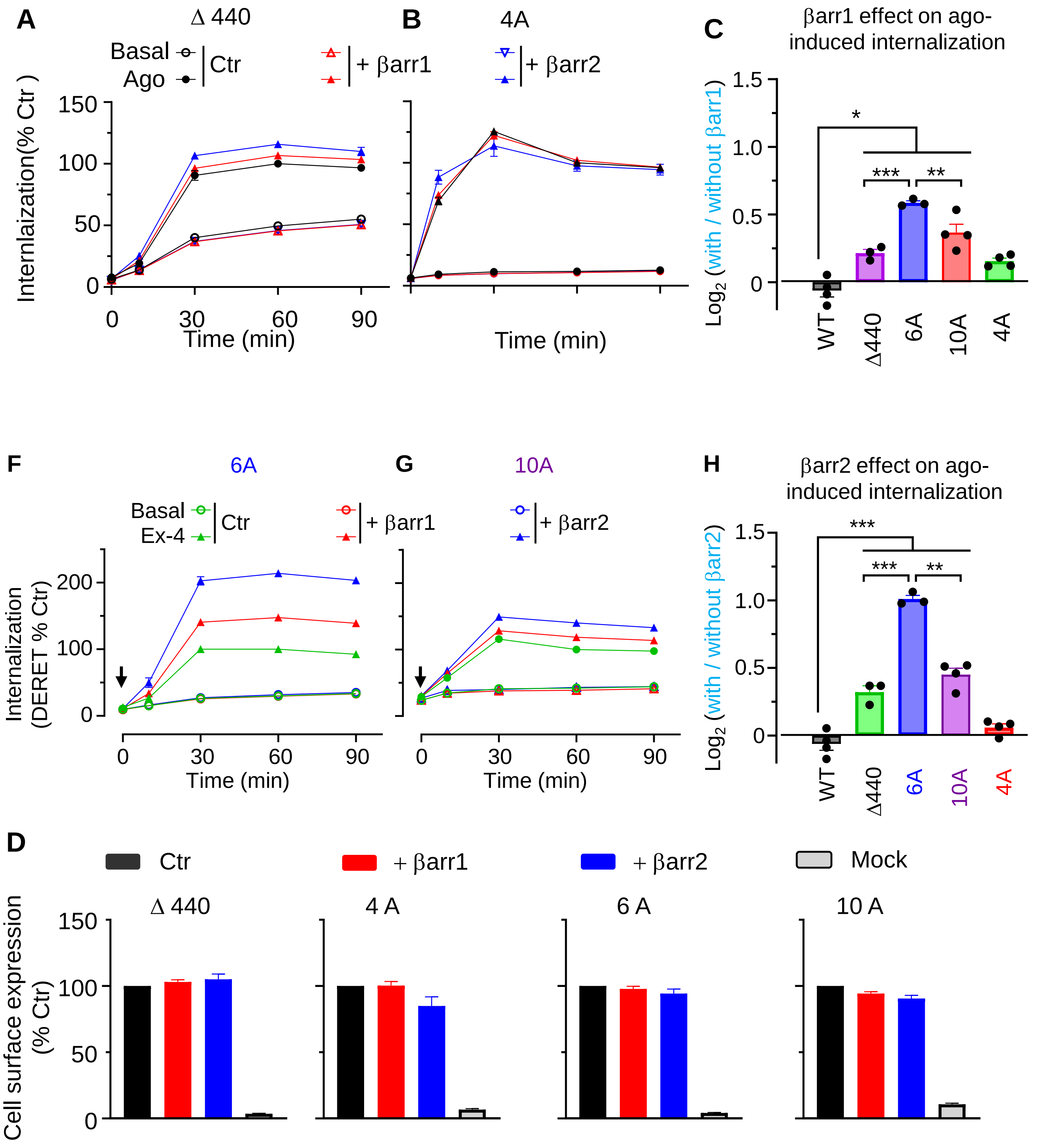

### Supp Fig 17. GRKs are involved in GLP-1R internalization.

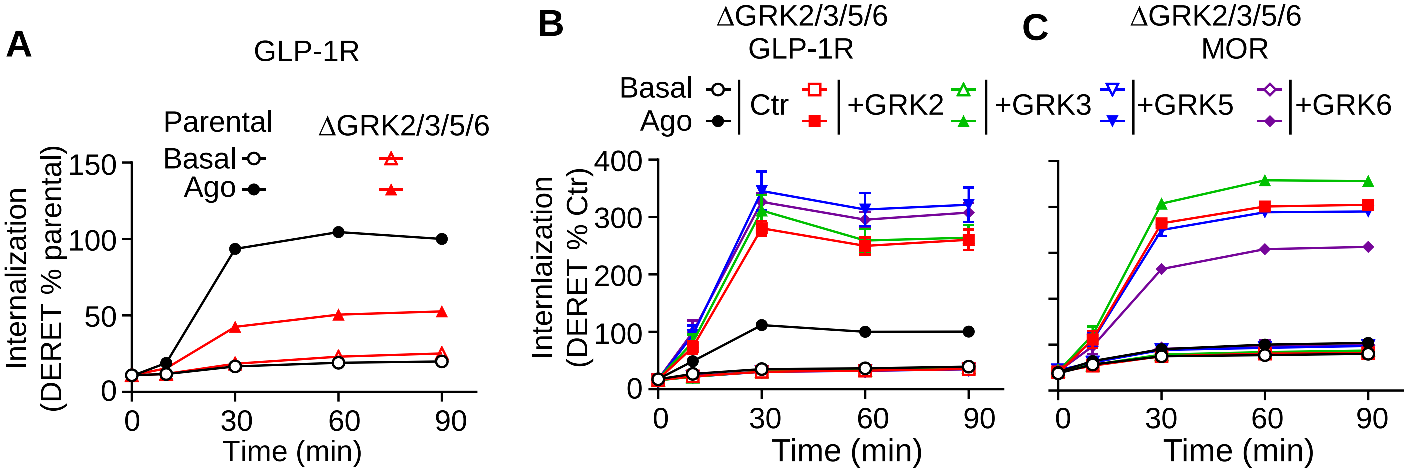

Supp Fig 18. Individual GRK effect on barr2 recruitment, AP2 recruitment, and the agonist-induced internalization of GLP-1R

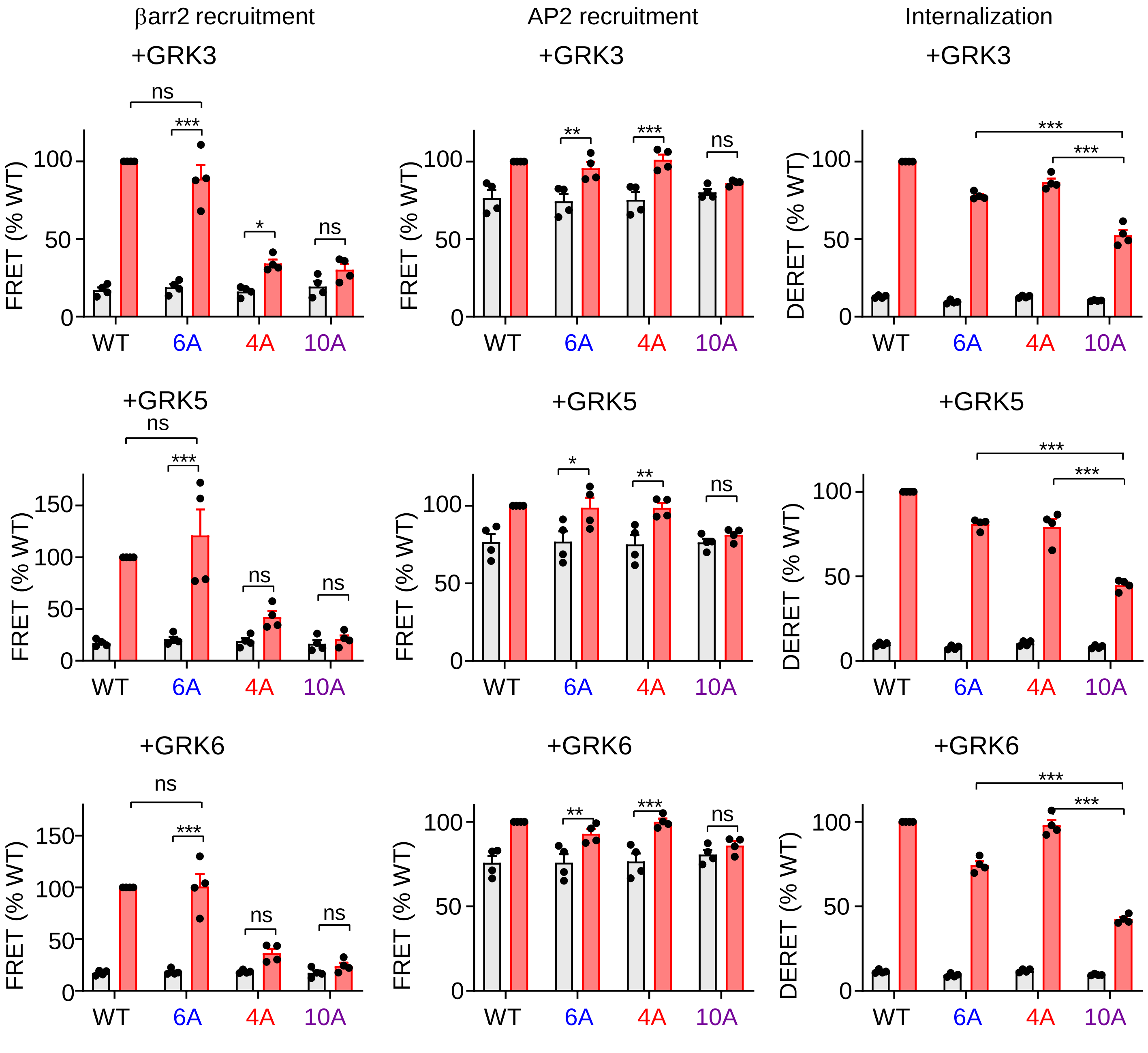

### Supp Fig 19. Expression and internalization profile of Natural genetic variants of GLP-1R

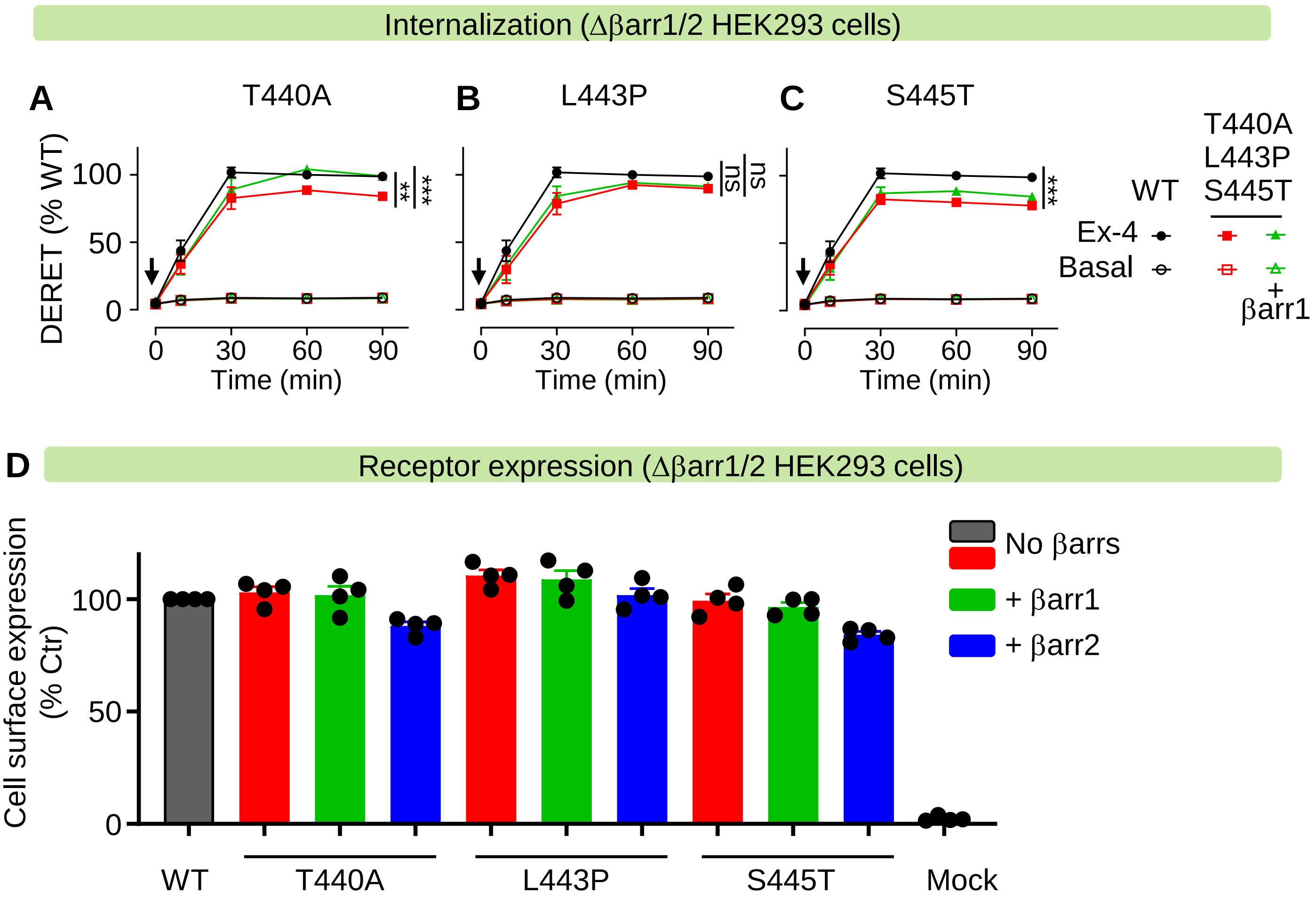
